# Success probability of high-affinity DNA aptamer generation by genetic alphabet expansion

**DOI:** 10.1101/2022.06.27.497860

**Authors:** Michiko Kimoto, Hui Pen Tan, Yaw Sing Tan, Nur Afiqah Binte Mohd Mislan, Ichiro Hirao

## Abstract

**Summary:** Nucleic acid aptamers as antibody alternatives bind specifically to target molecules. These aptamers are generated by isolating candidates from libraries with random sequence fragments, through an evolutionary engineering system. We recently reported a high-affinity DNA aptamer generation method that introduces unnatural bases (UBs) as a fifth letter into the library, by genetic alphabet expansion. By incorporating hydrophobic UBs, the affinities of DNA aptamers to target proteins are increased over 100-fold, as compared to those of conventional aptamers with only the natural four letters. However, there is still plenty of room for improvement of the methods for routinely generating high-affinity UB-containing DNA (UB-DNA) aptamers. The success probabilities of the high-affinity aptamer generation depend on the existence of the aptamer candidate sequences in the initial library. We estimated the success probabilities by analysing several UB-DNA aptamers that we generated, as examples. In addition, we investigated the possible improvement of conventional aptamer affinities by introducing one UB at specific positions. Our data revealed that UB-DNA aptamers adopt specific tertiary structures, in which many bases including UBs interact with target proteins for high affinity, suggesting the importance of the UB-DNA library design.

## 1. Introduction

Nucleic acid aptamers are single-stranded DNA or RNA molecules that bind specifically to a wide variety of targets, such as small molecules, sugars, proteins, and cells. Aptamers are generated by an evolutionary engineering method called SELEX (<u>S</u>ystematic <u>E</u>volution of <u>L</u>igands by <u>EX</u>ponential enrichment), through repetitive processes of selection and amplification using nucleic acid libraries with randomised sequences [1, 2]. Once the aptamer sequences have been determined by SELEX, the aptamers and their modifications are manufactured by solid-phase chemical synthesis and can be used as antibody alternatives. However, one main issue is the aptamers’ insufficient affinities to targets, because of the limited chemical diversity, especially the low hydrophobicity, of the nucleic acid components, A, G, C, and T/U, for interactions with hydrophobic regions of target proteins. Thus, the practical applications of nucleic acid aptamers are still limited, and many improved methods for modified aptamer generation have been reported [3-8]. Especially, the use of a chemically expanded library is widely considered to be advantageous for generating high-affinity aptamers, but a few studies have experimentally or theoretically confirmed the validity of this strategy [9].

Recently, we developed a novel DNA aptamer generation method called ExSELEX (genetic alphabet <u>Ex</u>pansion for <u>SELEX</u>), in which a hydrophobic unnatural base (UB), 7-(2-thienyl)-imidazo [4,5-b] pyridine (Ds), is introduced as a fifth letter [10-13] (Supplementary figure S1). The Ds-containing DNA (Ds-DNA) fragments can be amplified by PCR using the unnatural base pair (UBP) between Ds and 2-nitro-4-propynylpyrrole (Px) [14, 15] (figure 1a), allowing for Ds-DNA aptamer generation by ExSELEX. We generated several high-affinity Ds-DNA aptamers with 0.65–132 pM *K*D values, targeting vascular endothelial growth factor 165 (VEGF165), interferon γ (IFNγ) [10], von Willebrand Factor A1-domain (vWF) [16], and each serotype of dengue NS1 proteins [17] (figure 1b). However, the generation of Ds-DNA aptamers with high affinities (sub-nanomolar *K*D) is still a laborious and intricate process and low success probabilities, because of the higher complexity including the fifth letter than that in the four-letter DNA aptamer generation.

**Figure 1.**
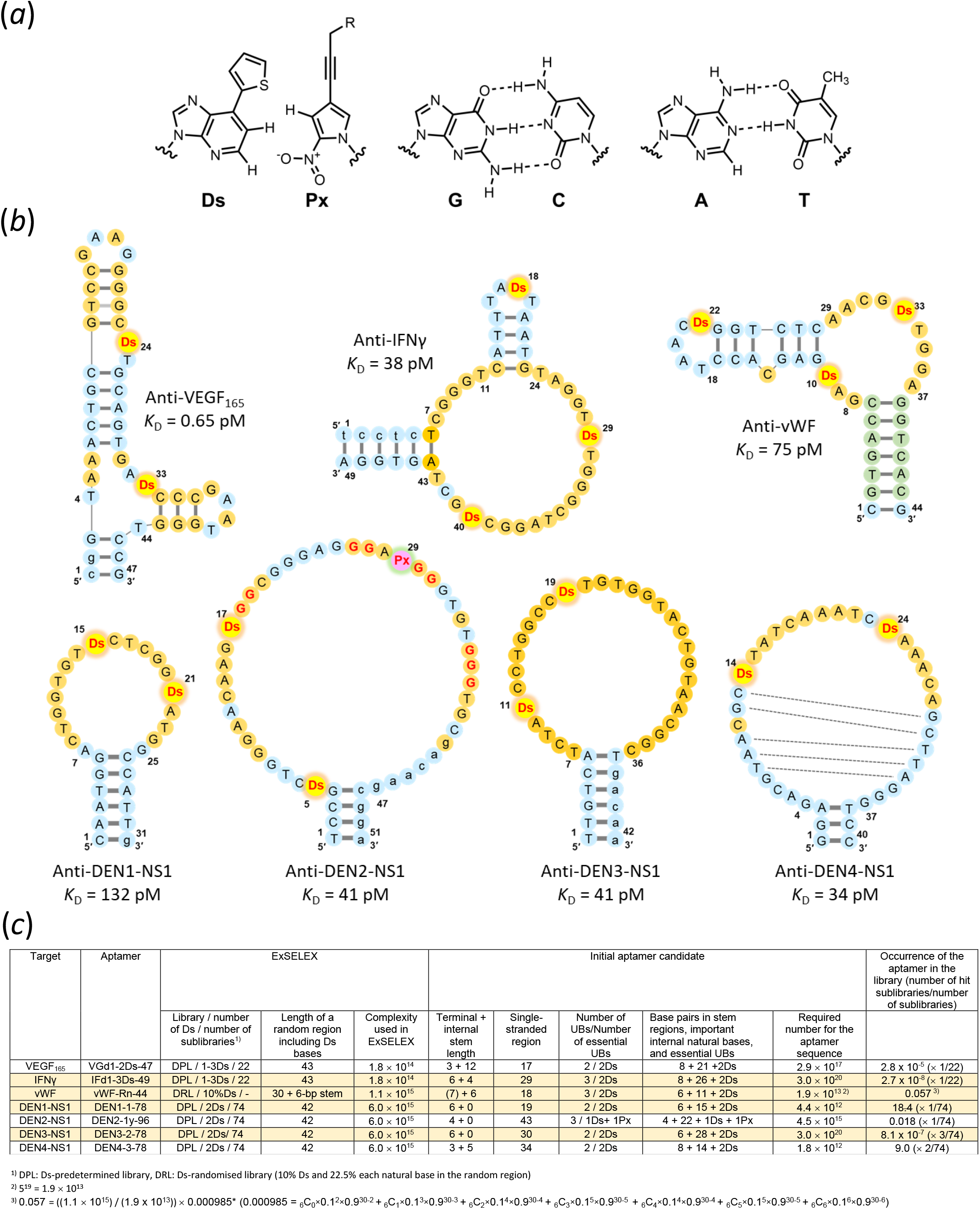
High-affinity Ds-DNA aptamer candidates obtained by ExSELEX. (a) Chemical structures of a hydrophobic unnatural base pair, Ds–Px, and the natural A–T and G–C base pairs. (b) Predicted secondary structures of high-affinity Ds-DNA aptamers targeting VEGF165, interferon γ (IFNγ), von Willebrand Factor A1-domain (vWF), and each serotype of dengue NS1 proteins. As for the anti-DEN1/2/3/4-NS1 aptamers, their core parts are shown by removing the flanking primer regions. The bases in the randomised and primer regions are shown in large and small letters, respectively. The important bases (orange circles) were estimated from the doped selections (anti-VEGF165 and anti-IFNγ aptamers) or the obtained sequence motifs (anti-DEN1-, DEN2-, and DEN4-NS1 aptamers). There is no information available for anti-vWF and DEN3-NS1, and thus the loop regions are tentatively estimated as the important bases. Green circles in the anti-vWF-aptamer are the original constant complementary sequences for stem formation. In the anti-DEN2-NS1 aptamer, the four G tract to form a G-quadruplex structure is shown by red Gs. The dotted lines in the anti-DEN4-NS1 aptamer represent a stem region found in the internal loop region. (c) Estimation of the occurrence of the aptamer candidates in the initial library based on ExSELEX data.

In the SELEX process, the complexity (number of different sequences) in the library is a key to increase the success probability of high-affinity aptamer generation, which relies on the existence of aptamer candidates in the initial library [7, 9, 18-23]. Greater complexity in the initial library increases the success probability of aptamer generation. However, a typical SELEX procedure limits the complexities to 1012–1015 different sequences, due to the scale of the handling volume and the concentration of DNA/RNA. For example, a representative modified RNA aptamer (*K*D = 10 pM), an initial precursor (V30.44) of pegaptanib [24], targeting VEGF165 was obtained from 1 nmol of a modified RNA library with a 30-base random region (N30), which contains 6.02 × 1014 sequences (10-9 × Avogadro’s constant) [25] (Supplementary figure S2). The aptamer consists of 19 internal stem-loop bases and a 4-base pair (bp) terminal stem. If each of the 19 bases in the internal stem-loop regions is essential or important and any base pairs are acceptable in the 4-bp terminal stem, then the occurrence of the 27-mer aptamer candidates would be (6.02 × 1014 × 12)/(7.0 × 1013 = 4(19+4)) in the library containing N30 and primer regions, suggesting that ∼103 aptamer candidate sequences were included in 1 nmol of the initial N30 library (refer to Supplementary figure S2 for the calculation).

In ExSELEX with the fifth letter, the complexity of the library becomes greater and more intricate than that of the conventional 4-letter libraries, theoretically reducing the success probability of high-affinity aptamer generation due to the scale limitations of the initial library. In general, the possible number of all the different sequences in the N30 library with the four natural letters is 430 = ∼1.15 × 1018. However, the number in the N30 library with the randomised five letters increases to 530 = ∼9.31 × 1020, indicating that only 1014–1015 different sequences (0.0001–0.001% of the total complexity) in the number are used in ExSELEX.

Here, we analysed the ExSELEX data for the high-affinity aptamers that we generated, to estimate the success probabilities of Ds-DNA aptamer generation from the viewpoint of five-letter DNA libraries. For the analysis, we determined the essential or important bases in the four Ds-DNA aptamers targeting IFNγ, vWF, and dengue virus serotypes 1 and 3 NS1 proteins by a gel-electrophoresis mobility shift assay (EMSA). We also examined the Ds incorporation into a conventional four-letter DNA aptamer, ARC1172, targeting vWF, to determine whether the hydrophobic UB improves the aptamer affinity and rationalised the data by structural modelling. These results reveal the importance of both the specific bases including Ds and the unique tertiary structures of each aptamer, providing information for further improvements in the design of UB-DNA libraries.

## 2. Materials and Methods

### (a) Materials

DNA aptamer variants, listed in Supplementary tables S1–S5, were chemically synthesised with an Oligonucleotide synthesiser nS-8 (Gene Design) or an H8 DNA/RNA Synthesiser (K&A Laborgerate), using phosphoramidite reagents for the natural and Ds bases. The Ds phosphoramidite was prepared as described previously [26], and the phosphoramidites for the natural bases were purchased from Glen Research. The synthesised DNA fragments were purified by denaturing gel electrophoresis before use. Recombinant proteins, human vWF A1 domain (vWF: amino acids 1238 to 1481), human IFNγ, and dengue virus serotype-1 NS1 (DEN1-NS1) and serotype-3 NS1 (DEN3-NS1), were purchased from U-Protein Express, Peprotech, and Native Antigen Company, respectively.

### (b) EMSA

The relative binding efficiency of each aptamer variant to the target protein was determined by EMSA, as described previously [16, 17, 27]. The conditions for the binding and gel electrophoresis, along with their respective buffers, are summarised in Supplementary table S6. The relative binding efficiency of each aptamer variant with the target protein was determined from the shifted band patterns on the gels, detected with a LAS-4000 bioimager (Fuji Film) after staining with SYBR Gold. The band densities of free and complexed DNAs were quantified using the Multigauge software (Fuji Film), and the relative binding (%) was calculated by normalisation with the shifted ratio for the original aptamer in the same gel.

### (c) Molecular dynamics (MD) simulations

To investigate the target interactions with some of the substituted variants of ARC1172, we performed molecular dynamics simulations. Details of the preparation of the starting structures, MD simulations, and binding free energy calculations are described in the supplementary information.

## 3. Results

### (a) Success probability of high-affinity aptamer generation by ExSELEX

First, we reviewed the success probability of the high-affinity Ds-DNA aptamer (*K*D < nM) generation by considering seven aptamers that we generated by ExSELEX, targeting VEGF165, IFNγ, vWF, and four dengue virus serotypes of NS1 proteins (DEN1-NS1 to DEN4-NS1) (figure 1b). These aptamers were isolated from single-stranded Ds-DNA libraries with 1014–1015 different sequences (complexities) (figure 1c), after 7–10 rounds of selection and PCR amplification. All of the isolated aptamer candidates with high affinities had specific sequences containing two or three Ds bases, flanked by complementary terminal stem regions (figure 1b).

In the anti-VEGF165, IFNγ, DEN1-NS1, DEN2-NS1, DEN3-NS1, and DEN4-NS1 aptamer generations, we used a mixture of Ds-predetermined sublibraries (DP library) (Supplementary figure S1c), in which one to three Ds bases are embedded at specific positions within a natural-base randomised region. For the anti-VEGF165 and IFNγ aptamer generation, 22 sublibraries (total complexity: 1.8 × 1014) with one to three Ds bases were used [10], and for each serotype of anti-DEN-NS1 aptamer generation, 74 sublibraries (total complexity: 6 × 1015) with two Ds bases were used [17]. In the anti-vWF aptamer generation, we employed a Ds-randomised (5-letter-randomised) library (DR library; complexity: 1.1 × 1015) consisting of 10% Ds and 22.5% each natural base in an N30 region flanked by conserved complementary 6-base sequences that form a stem, as well as PCR primer sequences [16] (Supplementary figure S1e).

The isolated anti-IFNγ and vWF aptamers contain three Ds bases. After the characterisation and optimisation of each aptamer, we determined that two Ds bases, at positions 29 and 40 in the anti-IFNγ aptamer and positions 10 and 33 in the anti-vWF aptamer, are essential for their high affinities (figure 1b). The non-essential Ds bases at position 18 for anti-IFNγ and position 22 for anti-vWF are each located in a small loop region, which can be replaced with a more stable mini-hairpin sequence (Supplementary figure S3) [16, 28, 29].

The anti-DEN2-NS1 aptamer contains one Px base at position 29, as well as two Ds bases. The Px base is the pairing partner of Ds for PCR amplification, and this Px base in the aptamer appeared by the misincorporation of Px opposite natural bases in templates during PCR in ExSELEX. This Px at position 29 and one Ds at position 17 are essential for high affinity binding, and the other Ds at position 5 is not essential. The anti-DEN2-NS1 aptamer forms a G-quadruplex structure with four G tracts shown by red Gs in figure 1b [17].

The success probability of each aptamer generation was estimated from the occurrence of the aptamer in the library, which was determined from the complexity in the initial library divided by the required number for the aptamer candidate sequence in each ExSELEX (figure 1c). The required number for aptamer candidate sequence, in which at least one aptamer sequence possibly appears, was determined using the number of essential/important bases and the stem length in each aptamer. The important bases are indicated in orange color circle in the secondary structures in figure 1b. Each important base was estimated by the data obtained from the second ExSELEX, using doped libraries for anti-VEGF165 and IFNγ aptamer generation, in which we chose >96% conservation bases [10], or from the conserved sequences in the isolated clones for anti-DEN-NS1 aptamer generation [17]. For example, in the anti-VEGF165 aptamer, there are 21 important natural bases including two stem regions, as well as 2 Ds bases, and 8 base pairs, in which any base pairs are acceptable, in the terminal and internal stem regions. Thus, the required number for the aptamer sequences is 4(21+8) = 2.9 × 1017. In the anti-DEN3-NS1 aptamer generation, only one sequence in the isolated clones was obtained, and no consensus sequences in the clone were identified. Thus, in the calculation of the anti-DEN3-NS1 aptamer, we used all of the bases (28 natural bases and 2 Ds bases) in the loop region and the 6-bp terminal stem, and the tentative required number for the aptamer sequence is 4(28+6) = 3.0 × 1020. From the sequencing data of the anti-DEN4-NS1 aptamer generation, we found stem regions in the internal loop region (figure 1b), and thus the aptamer contains eight base pairs (three in terminal and five in loop regions) in the secondary structure.

In the anti-vWF aptamer generation using the DR library, we determined the required number for the aptamer sequence by a different method. The regions of the terminal stem and the loop at positions 18–22 were ignored in the calculation, because the terminal stem was embedded into the initial library as a constant sequence, and the bases in the loop regions were not essential for tight binding. Since we had no data about the important bases in the other regions, all 13 bases (including two Ds bases) in the single-stranded region were treated as the important bases in the five letters, and two internal stem regions were treated as the stems with non-essential bases. Thus, the required number for the anti-vWF aptamer is 5(13+6) = 1.9 × 1013.

From these required numbers for each aptamer sequence and the complexities of each initial library used, we calculated the occurrence of the aptamer in the library, which is the theoretical number of aptamer candidate sequences found in the initial libraries (figure 1c). In ExSELEX using DP libraries, the complexity of the initial library was divided by the required numbers for the sequence and by the total sublibrary number used, and then multiplied by the number of the sublibraries that include two Ds bases at the same interval (-Ds-Nn-Ds-) and appropriate positions (“number of hit sublibraries” in figure 1c). For example, the occurrence of the aptamer in the library for the anti-DEN4-NS1 aptamer generation is ((6.0 × 1015) / (1.8 × 1013)) × 2/74 = ∼9.0, since two of the 74 sublibraries have two Ds bases at the same interval and appropriate positions with the aptamer. As for the anti-DEN2-NS1 aptamer, only one Ds base is essential, and thus the aptamer could possibly be isolated from all 74 sublibraries. In the anti-vWF aptamer generation, we calculated the occurrence of the aptamer in the library by a different method due to the use of the DR library (refer to figure 1c and Supplementary figure S4).

The occurrence of the aptamer candidates in the library (figure 1c) revealed that the library size used for most of the aptamer generations, except for the anti-DEN1-NS1, DEN2-NS1, and DEN4-NS1 aptamers, was too small. However, the aptamers were successfully obtained from the libraries, suggesting that some factors used for the calculation might not reflect that fact. One of the uncertainties in the calculation is the determination of the important or essential bases in each aptamer. Thus, we re-evaluated the important/essential bases in the single-stranded regions of some aptamers, focusing on the anti-IFNγ, vWF, DEN1-NS1, and DEN3-NS1 aptamers, by EMSA using a series of sequence variants for each aptamer.

For the assay, we chemically synthesised a variant set for each aptamer by point transition mutation (A↔ G, T↔C) in the single-stranded regions. We used the optimised sequences for some aptamers, as shown in Supplementary figures S3 and tables S1 to S4. In the optimisation, the terminal stem lengths were extended and the A–T pairs were replaced with G–C pairs in the stem of each aptamer [10, 12, 17, 29]. The small internal stem-loop sequences in the anti-IFNγ and vWF aptamers were replaced with thermally stable mini-hairpin sequences, CGCGAAGCG or CCGAAGG [16, 29] (Supplementary tables S1 and S2). Therefore, these two aptamers are highly stabilized thermally. Binding analyses of these anti-IFNγ and anti-vWF aptamers to their targets were performed using 25 nM aptamer variants and 50 nM target proteins, and the complexes were separated from the free DNA by electrophoresis on a gel containing 3 M urea at 30–37°C for EMSA. In our ExSELEX procedure, we had employed stringent washing conditions of the DNA-target complexes to isolate high-affinity aptamers. As a result, some aptamers had very high affinities (less than several ten pM of *K*D values) to their targets. Thus, we used the gel analysis conditions in the presence of 3 M urea for anti-IFNγ and anti-vWF aptamers and their variants to identify the differences between their nM and pM *K*D values [16] (figure 2a, Supplementary figure S5 and table S6). The binding analyses of anti-DEN1-NS1 and anti-DEN3-NS1 aptamers, which have no internal mini-hairpin sequences, were performed using 50 nM aptamer variants and 25 nM of each DEN-NS1 hexamer, and the complexes were detected on a native gel at 30°C, in the absence of urea (Supplementary figures S6 and S7, and table S6). The anti-DEN1-NS1 aptamer has an additional mini-hairpin sequence, CGCGAAGCG, at its 3’-terminus [17] (Supplementary table S3), since its target binding was slightly lower than that of the anti-DEN3-NS1 aptamer and the terminal stem stability affected its affinity (data not shown).

**Figure 2.**
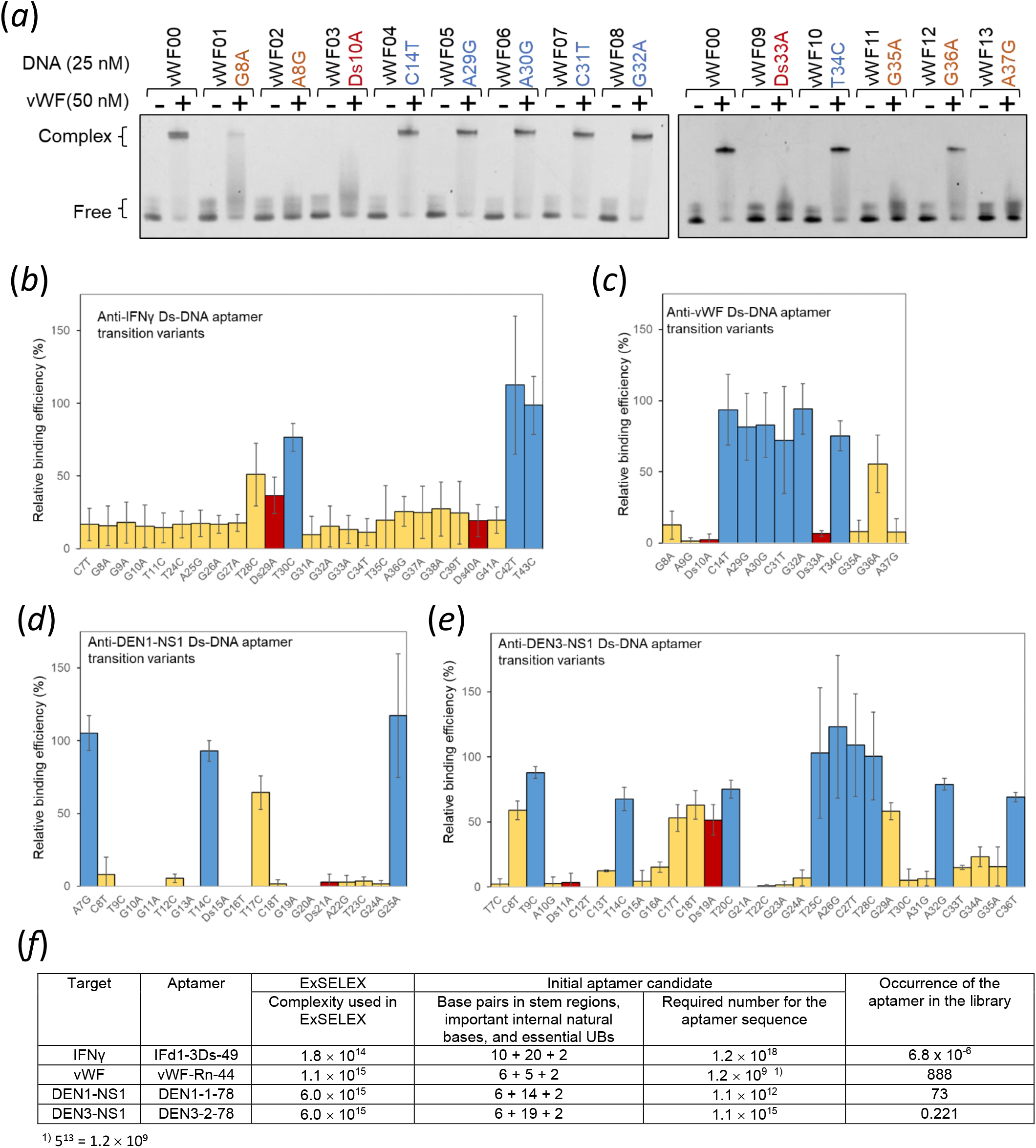
Binding analysis of Ds-DNA aptamer variants with point transition mutations. (a) EMSA for the anti-vWF Ds-DNA aptamer variants. (b–e) Relative binding efficiencies (%) of each variant of anti-IFNγ Ds-DNA aptamer (b), anti-vWF Ds-DNA aptamer (c), anti-DEN1-NS1 Ds-DNA aptamer, and (d) anti-DEN3-Ns1 Ds-DNA aptamer (e), determined with normalisation of the complex formation efficiency of each original Ds-DNA aptamer (Supplementary figure S3) from EMSA (Supplementary figures S5–S7). The Ds positions are shown by red bars. The positions that exhibited less than 65% of the relative binding efficiencies (average) were assigned as the essential base positions and are shown by orange bars. (f) Re-estimation of the occurrence of the aptamer candidates in the initial library used, according to the important/essential bases identified by EMSA.

The binding proportion (%) of the complex of each aptamer and its variants with target proteins was measured from the band densities of the complex and the unbound aptamer. The relative binding efficiencies of each aptamer variant relative to the original aptamer are summarised in figure 2b–2e. The binding efficiencies of most of the variants for each aptamer were lower than those of the original one, and some base mutations significantly reduced the binding to their targets. Each mutated base position that exhibited less than 65% of the relative binding efficiencies (average) was assigned as an important base position for high affinity (sub-nanomolar *K*D) in each aptamer, based on our previous EMSA results [27].

We re-calculated the required numbers for aptamer sequences and the occurrence of the aptamer in the library for the anti-IFNγ, vWF, DEN1-NS1, and DEN3-NS1 aptamers, using the important bases determined by the EMSA experiments (figure 2f). Since the numbers of essential or important bases obtained by EMSA are smaller than those in figure 1c, reasonable numbers of the anti-vWF, DEN1-NS1, and DEN3-NS1 aptamer sequences are included in the initial library. However, the occurrence of the anti-IFNγ aptamer in the library is still very low, and theoretically, a library with 1.2 × 1018 complexity would be required for this aptamer generation.

### (b) High-affinity aptamer generation by natural to unnatural base mutations in conventional aptamers

Another intriguing issue of UB-DNA aptamer generation is whether high-affinity Ds-DNA aptamers can be obtained by replacing any natural base with Ds in known 4-letter aptamers. As a model system, we chose the 41-mer DNA aptamer targeting vWF, ARC1172 (or ARC1779), which inhibits vWF binding to platelet-receptor glycoprotein Ibα and is under phase II clinical trials [30-33]. Based on the secondary structure of the aptamer and the tertiary structure in the complex with vWF [34] (figure 3a), we chemically synthesised two sets of its variants; variants with a point transition mutation and variants with the Ds-replacement of each base in the single-stranded regions (positions, 7, 8, 10, 21, 22, 27–31) (Supplementary table S5). The relative binding efficiencies (%) of each variant in the two sets were determined by EMSA, using 100 nM of each aptamer variant and 200 nM vWF (figure 3b, Supplementary figure S8 and table S6). The data revealed that the positions at T10 and G28 were acceptable as the Ds-replacement positions, while the original natural bases at positions 8, 10, 21, 22, and 27 are relatively important for the tight binding. However, the affinities of two variants, T10Ds and G28Ds, did not exceed that of the original aptamer.

**Figure 3.**
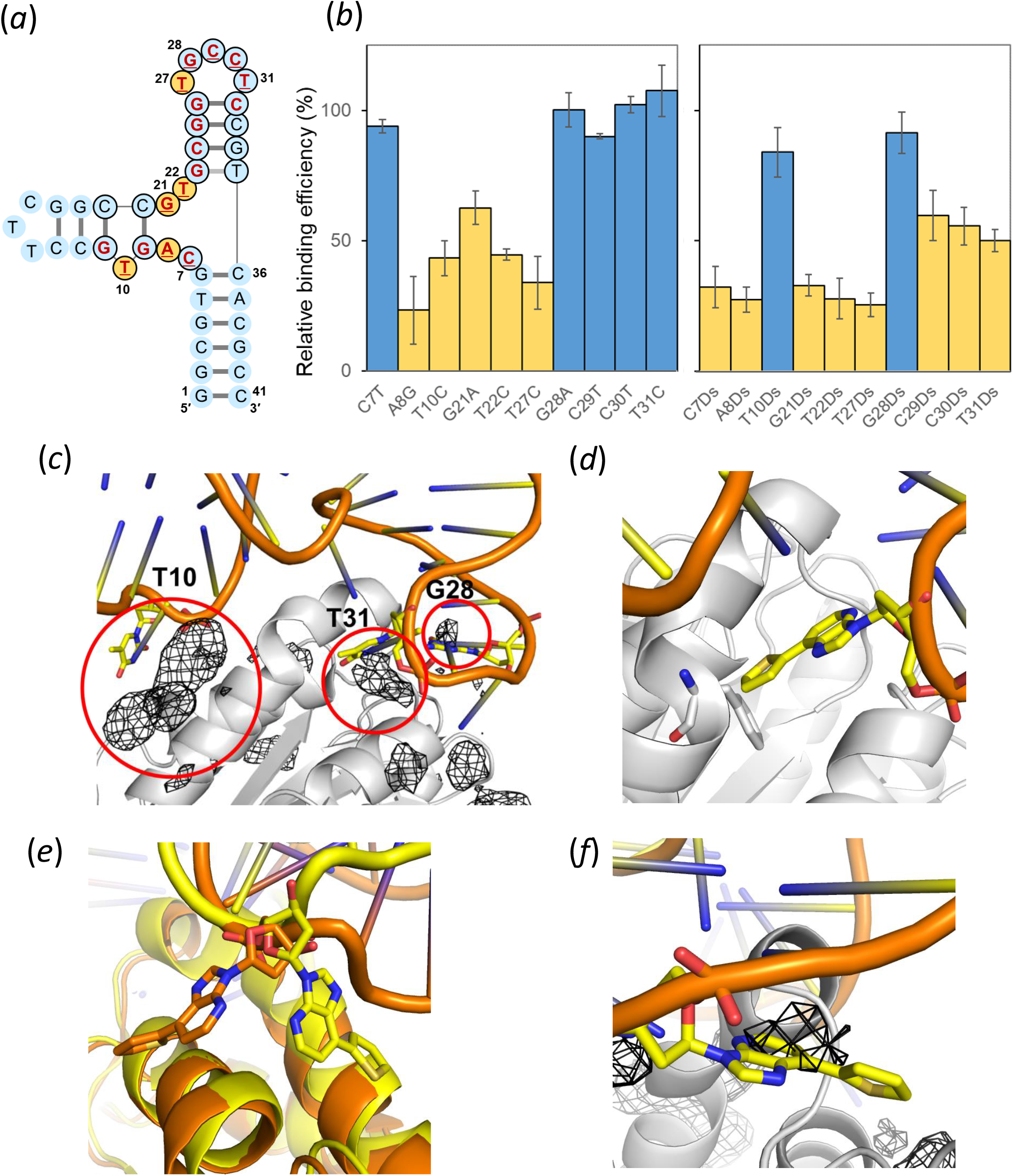
Binding of ARC1172 variants with vWF. (a) Schematic illustration of ARC1172. Nucleotides with bold red letters are within 4.2 Å of vWF in the complex [34]. The consensus nucleotides identified for high-affinity binding to vWF are indicated in black circles [34]. The 10 positions for point mutations (transition or Ds incorporation) are underlined. The essential bases assigned from the EMSA (Supplementary figure S8) are shown in orange circles. (b) Relative binding efficiencies (%) of ARC1172 variants with vWF. The positions that exhibited less than 65% of the relative binding efficiencies (average) were assigned as the essential base positions and shown by orange bars. (c) Binding sites detected near the aptamer-vWF binding interface. Benzene occupancy maps (black mesh) overlaid on the structure of ARC1172 complexed with vWF (PDB 3HXQ), with detected binding sites near the aptamer-vWF binding interface circled in red. vWF is shown in white, and the aptamer is shown in yellow and orange. (d) Modelled structure of the vWF–ARC1172 (T31Ds) complex, showing steric clash between Ds31 (yellow) and Phe1397 (white). (e) MD snapshot of the vWF–ARC1172 (T10Ds) complex, showing alternative conformations of Ds10 (yellow and orange). (f) MD snapshot of the vWF–ARC1172 (G28Ds) complex, showing the overlap of Ds28 with the benzene map densities (black mesh).

To simulate the potential binding sites for Ds in vWF, we first performed ligand-mapping molecular dynamics (LMMD) simulations. LMMD is a computational pocket detection method that uses small organic molecules called fragments to rapidly identify binding pockets in MD simulations [35-38]. Since the Ds nucleotide is highly hydrophobic, benzene was used as the probe of choice in the simulations.

Based on the benzene occupancy maps generated from LMMD simulations of vWF, we identified three hydrophobic binding sites near the aptamer binding interface. These hydrophobic binding sites are close to the sites where T10, G28 and T31 of ARC1172 bind (figure 3c), and positions T10 and G28 are consistent with the data from the Ds-replacement experiments. The results support that the Ds base interacts with a hydrophobic region in vWF. Position T10 corresponds to the binding site of Leu110 in botrocetin, a snake venom protein that binds to vWF and promotes dysfunctional platelet aggregation (Supplementary figure S9a), while position T31 corresponds to the binding site of Tyr291 in botrocetin and Tyr316 of the anti-vWF antibody NMC-4 (Supplementary figure S9b and S9c). The significant decline in the relative binding efficiency of the T31Ds variant might be attributed to steric clashes introduced by the bulkier nitrogenous base of Ds (figure 3d), while C31, being a pyrimidine base like thymine, is still acceptable for the binding as shown in figure 3b.

To understand the tolerance at positions T10 and G28 for the Ds replacement, we next performed MD simulations of vWF complexed with T10Ds and G28Ds variants. Since the T10 and G28 bases are solvent-exposed, the perturbation of the tertiary structures by the Ds replacement is likely to be minor. As expected, both variants remained bound to vWF throughout the simulations with little deviation from the initial conformation. The calculated binding free energies of the Ds-replaced variants are not significantly different from that of ARC1172 (Supplementary table S7). The 1-deazapurine and thiophene moieties of T10Ds alternated between two flipped conformations during the simulations, suggesting that the thiophene ring neither interacts strongly nor enhances vWF binding significantly (figure 3e). As a comparison, the hydrophobic interaction of the thiophene ring of the G28Ds variant only partially compensated the loss of two hydrogen bonding between G28 and Glu1389 in the original aptamer, hence resulting in its moderate affinity (figure 3f).

For further understanding, we also analysed one of the low-affinity variants, T27C, by MD simulations. In ARC1172, T27 forms hydrogen bonds with the amide oxygen and backbone nitrogen of Gln1391, which allows the adjacent G26 to move closer to vWF and interact with the backbone nitrogen of Arg1392 (Supplementary figure S10a). These hydrogen bonds are maintained in the MD simulations (figure S10b). However, thymine has reversed positions of hydrogen bond-donating and -accepting atoms from cytosine. This leads to electrostatic repulsion between C27 and Gln1391. MD simulations of the T27C variant complexed with vWF show that the DNA segment from G26 to C27 cannot approach as close to vWF as that of ARC1172. C27 moves away from vWF and instead only forms hydrogen bonds with the amide side chain of Gln1391. This results in a loss of hydrogen bonding interactions between G26 and vWF (Supplementary figure S10c). The computed average binding free energy of the T27C variant is less negative than that of ARC1172 (Supplementary table S7), which agrees with the EMSA experiments. The SPR analysis of the original aptamer and T27C variant also showed that the transition mutation T27C reduced the affinity to vWF (Supplementary figure S10d).

We also checked the root mean square deviation (RMSD) for each MD run, focusing on the Cα backbone of vWF in LMMD, as well as the Cα and DNA backbone atoms in the complexes of vWF with ARC1172, the T10Ds variant, the G28Ds variant, and the T27C variant (Supplementary figures S11 and S12). The RMSD plots confirmed that each MD simulation reached a well-equilibrated steady state.

## 4. Discussion

The research of UB-DNA aptamer generation methods has just barely started, and only two types of UBPs, Ds–Px and Z–P, have been used [10, 13, 39-41]. The additional letters significantly increase the chemical diversity of libraries, which augment the aptamers’ affinities but reduce the success probabilities of UB-DNA aptamer generation. Here, from the viewpoint of the library size and complexity, we estimated the success probabilities of the Ds-DNA aptamers that we have generated so far. The results, obtained by considering the important or essential bases and stem structures in each aptamer, revealed that the isolated aptamer sequences targeting vWF and the series of DEN-NS1 proteins possibly existed in their initial libraries, and these high-affinity aptamers were reasonably generated. The anti-DEN2-NS1 aptamer contains one Px base, which was accidentally misincorporated by mutation during the ExSELEX procedure, and thus it was a serendipitous and fortunate result. In addition, a mutation occurring during the PCR amplification step plays an important role to increase the sequence diversity of a library with limited complexity [10, 39].

Our results for the anti-IFNγ and anti-VEGF165 aptamers indicated the difficulty of generating these aptamers, and a library with greater complexity (∼1018) might be required. These two aptamer generations were our first success. It must have been beginner’s luck because we later improved the DP library for the anti-DEN-NS1 aptamer generation, increasing the occurrence of each anti-DEN-NS1 aptamer generation in the library, which contains only two essential Ds bases at every position in the natural-base randomised region.

From this aspect, we should more rationally investigate why these anti-IFNγ and anti-VEGF165 aptamers were generated against incredible odds. Most of the Ds-DNA aptamers contain two essential Ds bases, which are sufficient to exhibit high affinities to their targets (sub-nanomolar *K*D values). These two Ds bases in the five aptamers are separated by 5–10 natural bases. Some, but not all, of the natural bases are important for the tight binding. Our LMMD simulations demonstrated that the highly hydrophobic Ds base interacts with a hydrophobic site in target proteins. Since the Ds bases significantly contribute the tight binding to targets, one plausible speculation is that at first, two Ds bases interact with two hydrophobic sites on the target protein, and then surrounding natural base sequences interact or fit with the target surface areas. In this stage, several different natural base sequence contexts might be available for the tight binding. In the RNA aptamer generation targeting MS2 coat proteins, a different sequence from the native MS2 RNA that binds to its coat proteins was isolated, and the X-ray crystallography of the complex revealed that the binding features of the RNA aptamer were very similar to those of the native MS2 RNA even with the different sequence context and the secondary structures [42]. Therefore, several sequence variations of the natural bases between or around the two Ds bases could be employed for high affinity binding, and we successfully isolated one of such sequences in the anti-IFNγ and anti-VEGF165 aptamer generation, even from the DP library with limited complexity.

The two Ds and specific natural bases might form a unique tertiary structure to fit the shape of each target protein. The terminal stem that always appears in the high-affinity Ds-DNA aptamers could stabilise the tertiary structure. Our Ds replacement experiments for ARC1172 did not show any further improvement of the aptamer affinity by only a replacement of the natural base with Ds at the appropriate positions, indicating the importance of suitable natural base sequence contexts around the Ds bases, which were obtained from the library by ExSELEX. The combination of two Ds and some natural bases in a specific structure is important for high-affinity UB-DNA aptamer generation, and thus the significant improvement of natural-base-DNA aptamers that have already been generated, only by replacing some natural bases with Ds, can be difficult as shown in the anti-vWF Ds-DNA aptamer generation.

We have two types of libraries, DP and DR, and most of the high-affinity UB-DNA aptamers were obtained by ExSELEX using the DP libraries. Our representative aptamer generated by using the DR library is the anti-vWF aptamer. Since two Ds bases are sufficient for high-affinity DNA aptamer generation, DP libraries with two Ds bases are more suitable than DR libraries. In our 42-mer DP library containing two Ds bases for anti-DEN-NS1 aptamer generation, the total possible number with different sequence contexts is 440 = ∼1.2 × 1024 in each sublibrary. In contrast, the number in a 42-mer DR library is increased to 542 = ∼2.3 × 1029, reducing the success probability of aptamer generation. Furthermore, in the N30 library with 10% Ds bases used for the anti-vWF aptamer generation, only 22.8% of the sequences contained two Ds bases (Supplementary figure S4), and the other sequences containing more than four or five Ds bases reduced the PCR amplification efficiencies in ExSELEX. Even so, the anti-vWF aptamer generation succeeded, because only six natural bases in the single-stranded region of the aptamer were important for the tight binding, and 888 aptamer candidate sequences existed in the initial DR library (figure 2f). On the other hand, one of the strong points of the DR library is that the terminal stem sequences can be embedded within the library, as shown in the N30 DR library for anti-vWF aptamer generation [16] (Supplementary figure S1e). To incorporate the terminal stem sequence into the DP library, we must prepare many sublibraries for all combinations of two-Ds-base contexts in the single-stranded region of the library.

As one of the limitations of this study using the doped libraries and the transition mutation variants, note that our current estimation to identify the important bases supposes a simplified situation, including a degree of uncertainty. For example, in anti-IFNγ and anti-VEGF165 aptamer generation, we used the results obtained by a second ExSELEX with the doped library to determine the essential bases and stem regions in the aptamer. However, such a library does not include hit sequences with random insertions or deletions. The doped selection of the anti-VEGF165 aptamer revealed that the two loop sequences, **<u>G</u>**A**<u>A</u>**G (positions 16–19) and **<u>G</u>**A**<u>A</u>**T (positions 37–40), contain two important bases (underlined and in bold), but another analysis using non-essential nucleotide-deletion variants (position 19 or/and 40) showed the loss of the binding affinity [28]. Although this might be a special case, in which a stable mini-hairpin sequence appeared by this deletion, causing loss of the affinity, this shows that not all deletions of non-essential bases are acceptable. In our analysis using mutation variants, we employed only transition mutations to simplify the method and did not strictly examine all of the variants with other mutations including transversions. In addition, we assumed that any stem sequences are acceptable, but some stem sequences complementary to a loop region could cause substantial changes in the proper aptamer folding and affect the affinity to targets.

This study illustrates some potentially useful considerations during aptamer generation or optimisation. The design of DP and DR libraries for ExSELEX might be further improved. Using the results of LMMD simulation of potential UB (Ds) binding site of a target protein (vWF), more appropriate DP sublibrary set can be chosen, without expanding excessive resources. During optimisation of a high-affinity DNA aptamer, the method of screening its transition mutation variants using EMSA offers a simple method to identify the important bases as as an initial screening.

## Acknowledgments

We thank Ms Yurina Saiki for assistance with some of the EMSA experiments for ARC 1172. This work was supported by the Institute of Bioengineering and Bioimaging and the Bioinformatics Institute (Biomedical Research Council, Agency for Science, Technology and Research, Singapore) and the 2021 Horizontal Technology Programme Office Seed Fund at A*STAR (C-21-13-18-002).

## Authors’ Contributions

M.K. and I.H. conceived the study. M.K., H.P.T, Y.S.T., N.A.B.M.M., and I.H. contributed and analysed data. M.K., Y.S.T. and I.H. prepared the manuscript.

