## Supplementary Information for "Success probability of high-affinity DNA aptamer generation by genetic alphabet expansion"

**Supplementary figure S1. A method to generate unnatural-base DNA aptamers using ExSELEX**

**Supplementary figure S2. Example to estimate the occurrence of aptamer candidates for an initial precursor**

**Supplementary figure S3. Original Ds-DNA aptamers for the point transition mutation experiments**

**Supplementary figure S4. Probability calculation for the vWF-Rn-44 sequence contexts using the N30 library**

**Supplementary figure S5. EMSA for the anti-IFNγ Ds-DNA aptamer variants**

**Supplementary figure S6. EMSA for the anti-DEN1-NS1 Ds-DNA aptamer variants**

**Supplementary figure S7. EMSA for the anti-DEN3-NS1 Ds-DNA aptamer variants**

**Supplementary figure S8. EMSA for the anti-vWF ARC1172 (ARC00) aptamer variants**

**Supplementary figure S9. Hydrophobic binding sites detected near the aptamer–vWF binding interface**

**Supplementary figure S10. Binding modes of ARC1172 and its T27C variant with vWF**

**Supplementary figure S11. Cα root mean square deviations (RMSDs) for 10 independent ligand-mapping trajectories of vWF**

**Supplementary figure S12. RMSDs for the Cα and DNA backbone atoms in the complexes of vWF with (A, B) ARC1172, (C, D) T10Ds, (E, F) G28Ds, and (G, H) T27C**

**Supplementary table S1. Anti-IFNγ aptamer (48-mer) and its variants for the transition mutation experiments**

**Supplementary table S2. Anti-vWF aptamer (42-mer) and its variants for the transition mutation experiments**

**Supplementary table S3. Anti-DENV1-NS1 aptamer AptD1 (48-mer) and its variants for the transition mutation experiments**

**Supplementary table S4. Anti-DENV3-NS1 aptamer (50-mer) and its variants for the transition mutation experiments**

**Supplementary table S5. Anti-vWF aptamer ARC1172 (41-mer) and its variants for the transition mutation and Ds-replacement experiments**

**Supplementary table S6. Experimental conditions for EMSA**

**Supplementary table S7. Computed average binding free energy of ARC1172 aptamer and its variants**

**Methods**

**Supplementary figure S1.** A method to generate unnatural-base DNA aptamers using ExSELEX (genetic alphabet Expansion for SELEX). (a) Chemical structures of natural A–T and G–C base pairs and a hydrophobic unnatural base pair, Ds–Px. (b) ExSELEX to generate a Ds-containing DNA (Ds-DNA) aptamer using the Ds-DNA library and PCR involving the Ds-Px pair as a third pair. (c–e) Types of Ds-DNA libraries are categorised into a Ds-Predetermined Library (DP Library), as a mixture of sublibraries, each containing Ds bases at predetermined positions in the natural-base randomised region (c) and a Ds-Randomised Library (DR Library) without and with specific complementary sequences for stem formation (d and e).

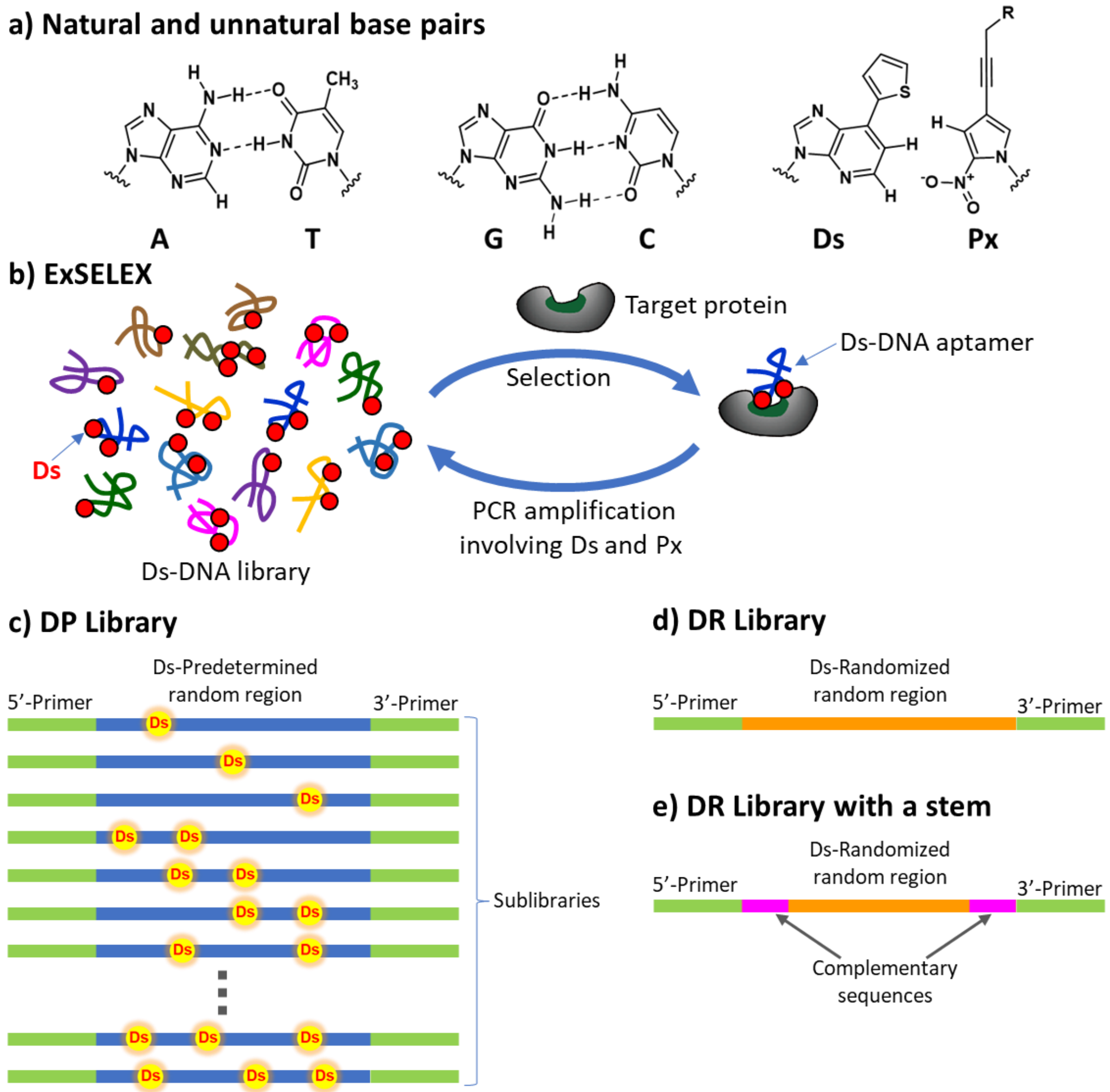

**Supplementary figure S2.** Example to estimate the occurrence of aptamer candidates for an initial precursor, VT30.44 (27-mer), of pegatanib in 1 nmol of the initial 30-base random (N30) library.

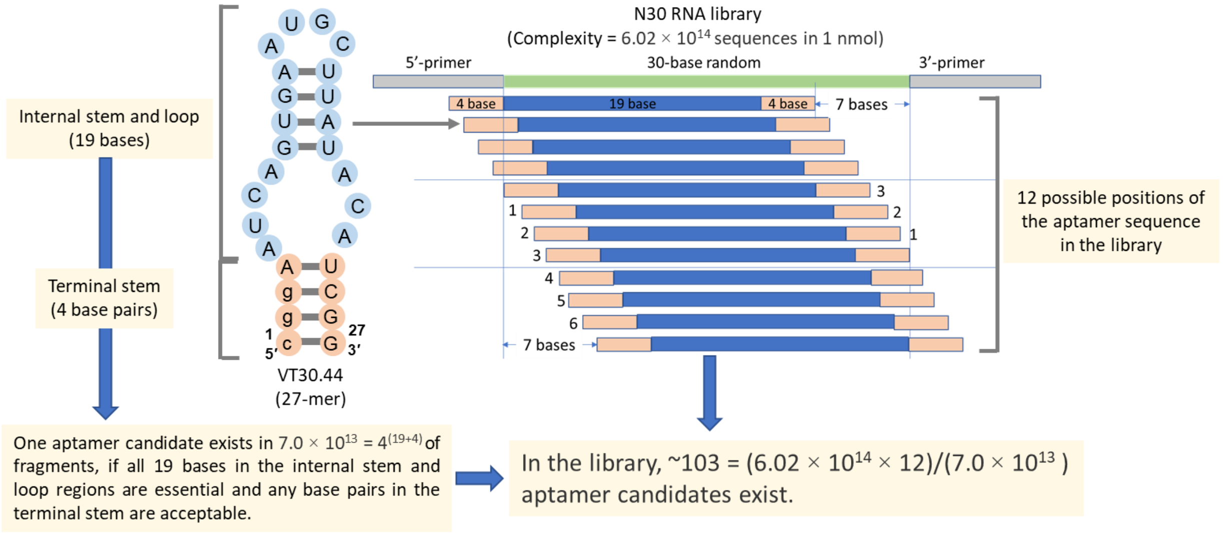


**Supplementary figure S3.** Original Ds-DNA aptamers for the point transition mutation experiments. The numbering of each aptamer sequence corresponds to the manner in figure 1b. The bases in orange circles are assigned as important bases by EMSA, and are different from those in figure 1b. The bases in pink circles are replaced to stabilise the aptamers for optimisation.

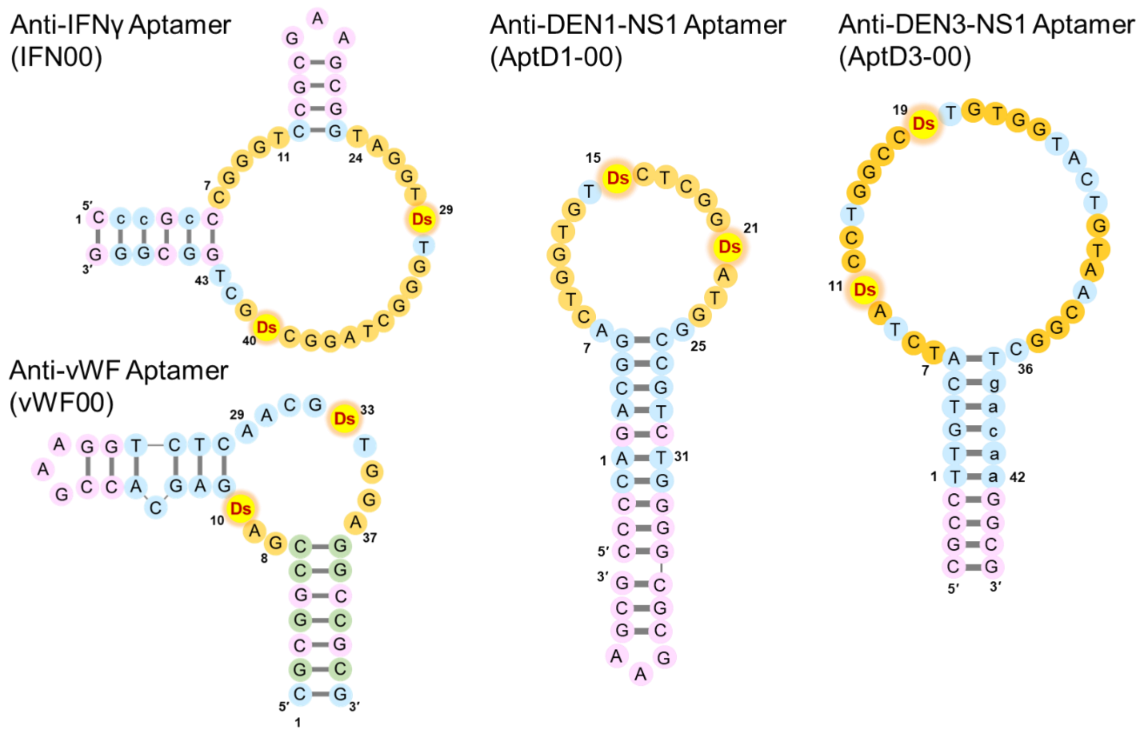

**Supplementary figure S4.** Probability calculation for the vWF-Rn-44 sequence contexts using the N30 library with Ds at 10% and each natural base at 22.5%.

| Numer of Ds (n) | Variations in sequence | Probability of the sequence variation | | cc(Ds)bbbcbbbnnnnnnbbbbbbcccc(Ds)cccc | | | |
|---|---|---|---|---|---|---|---|
| | $_{n}C_{30}\times4^{(30-n)}$ | Ds 20% & nautral base: 20% each $_{n}C_{30}\times0.2^{n}\times0.8^{(30-n)}$ | Ds 10% & nautral base: 22.5% each $_{n}C_{30}\times0.1^{n}\times0.9^{(30-n)}$ | $_{6}C_{x}$ | x = | $6Cx\times0.1^{n}\times0.9^{(30-n)}$ | Probability |
| 0 | 1.2E+18 | 1.24E-03 | 4.24E-02 | | | | |
| 1 | 8.6E+18 | 9.28E-03 | 1.41E-01 | | | | |
| **2** | 3.1E+19 | **3.37E-02** | **2.28E-01** | 1 | 0 | 0.000523348 | |
| **3** | 7.3E+19 | 7.85E-02 | 2.36E-01 | 6 | 1 | 0.000348898 | |
| **4** | 1.2E+20 | 1.33E-01 | 1.77E-01 | 15 | 2 | 9.69162E-05 | **0.000985** |
| **5** | 1.6E+20 | 1.72E-01 | 1.02E-01 | 20 | 3 | 1.4358E-05 | |
| **6** | 1.7E+20 | 1.79E-01 | 4.74E-02 | 15 | 4 | 1.1965E-06 | |
| **7** | 1.4E+20 | 1.54E-01 | 1.80E-02 | 6 | 5 | 5.31776E-08 | |
| **8** | 1.0E+20 | 1.11E-01 | 5.76E-03 | 1 | 6 | 9.84771E-10 | |
| 9 | 6.3E+19 | 6.76E-02 | 1.57E-03 | | | | |
| 10 | 3.3E+19 | 3.55E-02 | 3.65E-04 | | | | |
| 11 | 1.5E+19 | 1.61E-02 | 7.38E-05 | | | | |
| 12 | 5.9E+18 | 6.38E-03 | 1.30E-05 | | | | |
| 13 | 2.1E+18 | 2.21E-03 | 2.00E-06 | | | | |
| 14 | 6.2E+17 | 6.71E-04 | 2.69E-07 | | | | |
| 15 | 1.7E+17 | 1.79E-04 | 3.19E-08 | | | | |
| 16 | 3.9E+16 | 4.19E-05 | 3.33E-09 | | | | |
| 17 | 8.0E+15 | 8.63E-06 | 3.04E-10 | | | | |
| 18 | 1.5E+15 | 1.56E-06 | 2.44E-11 | | | | |
| 19 | 2.3E+14 | 2.46E-07 | 1.71E-12 | | | | |
| 20 | 3.2E+13 | 3.38E-08 | 1.05E-13 | | | | |
| 21 | 3.8E+12 | 4.03E-09 | 5.54E-15 | | | | |
| 22 | 3.8E+11 | 4.12E-10 | 2.52E-16 | | | | |
| 23 | 3.3E+10 | 3.58E-11 | 9.74E-18 | | | | |
| 24 | 2.4E+09 | 2.61E-12 | 3.16E-19 | | | | |
| 25 | 1.5E+08 | 1.57E-13 | 8.41E-21 | | | | |
| 26 | 7.0E+06 | 7.53E-15 | 1.80E-22 | | | | |
| 27 | 2.6E+05 | 2.79E-16 | 2.96E-24 | | | | |
| 28 | 7.0E+03 | 7.47E-18 | 3.52E-26 | | | | |
| 29 | 1.2E+02 | 1.29E-19 | 2.70E-28 | | | | |
| 30 | 1.0E+00 | 1.07E-21 | 1.00E-30 | | | | |
| **Total** | **9.3E+20** | **1.0E+00** | **1.0E+00** | | | | |

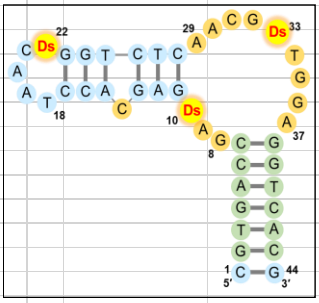

**Supplementary figure S5.** EMSA for the anti-IFNγ Ds-DNA aptamer variants.

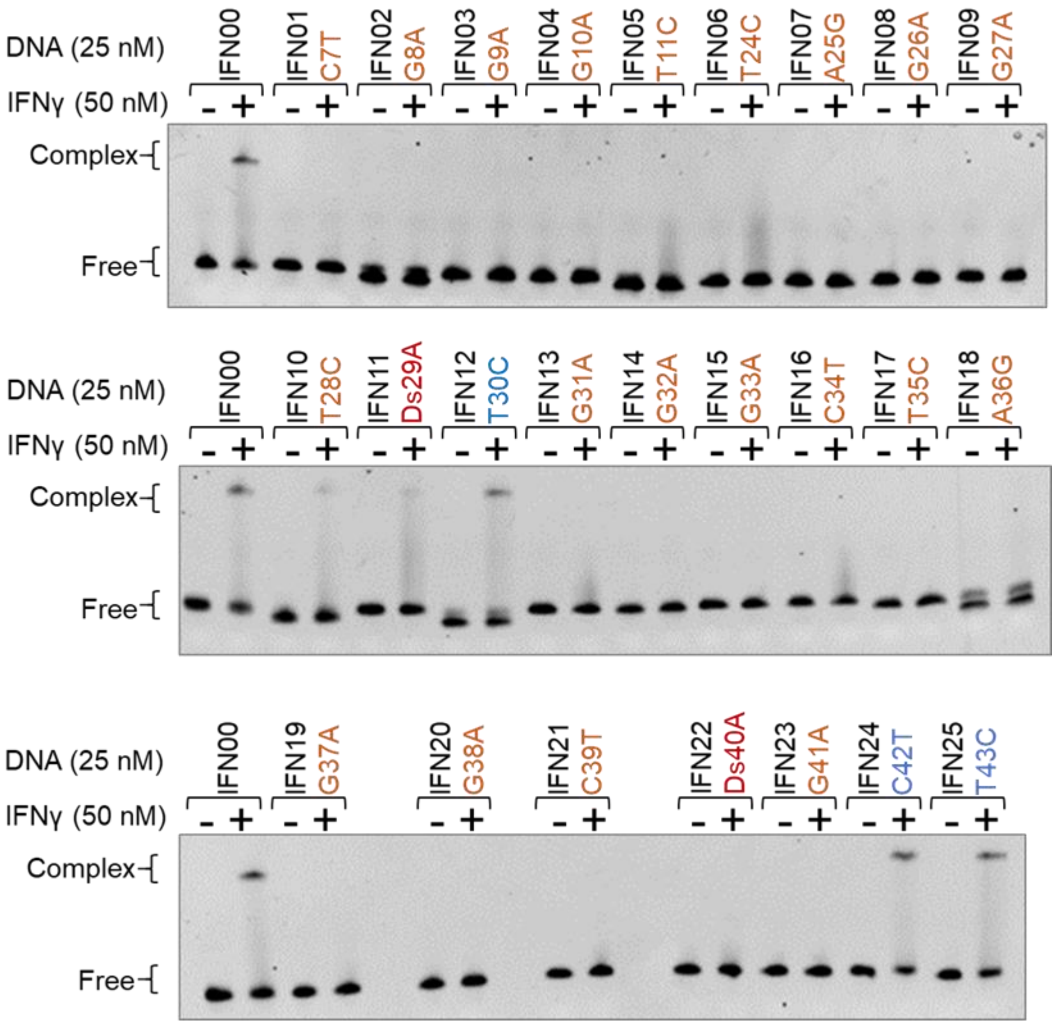


**Supplementary figure S6.** EMSA for the anti-DEN1-NS1 Ds-DNA aptamer variants.

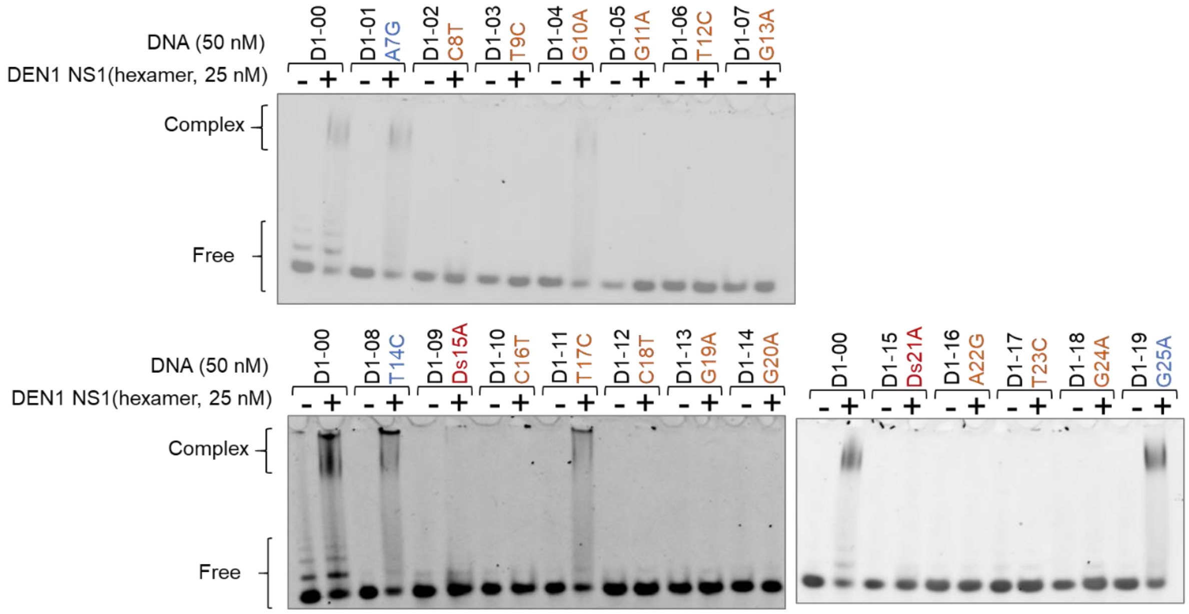

DNA (50 nM)
DEN1 NS1(hexamer, 25 nM)
D1-00
D1-01 A7G
D1-02 C8T
D1-03 T9C
D1-04 G10A
D1-05 G11A
D1-06 T12C
D1-07 G13A
Complex
Free
DNA (50 nM)
DEN1 NS1(hexamer, 25 nM)
D1-00
D1-08 T14C
D1-09 Ds15A
D1-10 C16T
D1-11 T17C
D1-12 C18T
D1-13 G19A
D1-14 G20A
D1-00
D1-15 Ds21A
D1-16 A22G
D1-17 T23C
D1-18 G24A
D1-19 G25A
Complex
Free

**Supplementary figure S7.** EMSA for the anti-DEN3-NS1 Ds-DNA aptamer variants.

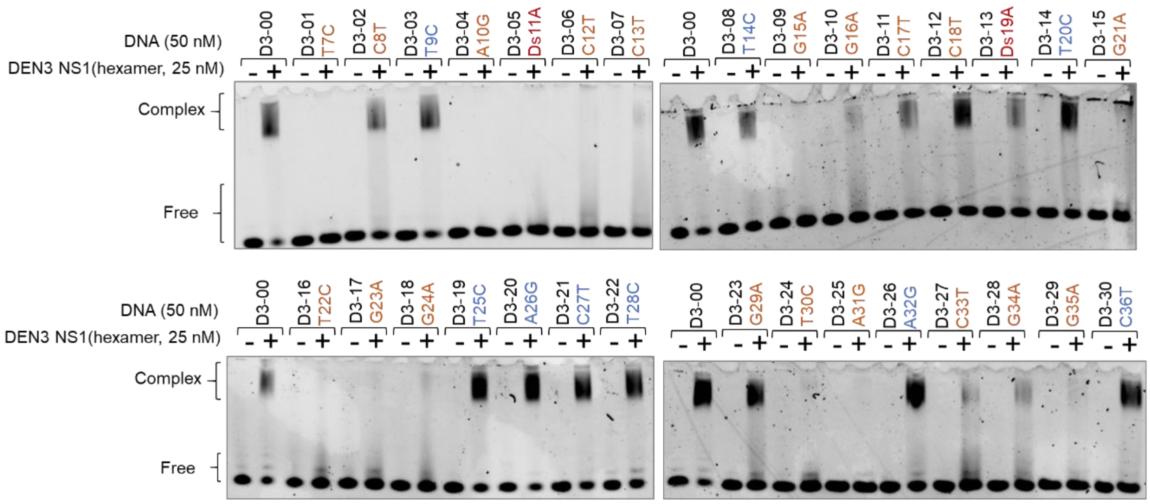


**Supplementary figure S8.** EMSA for the anti-vWF ARC1172 (ARC00) aptamer variants.

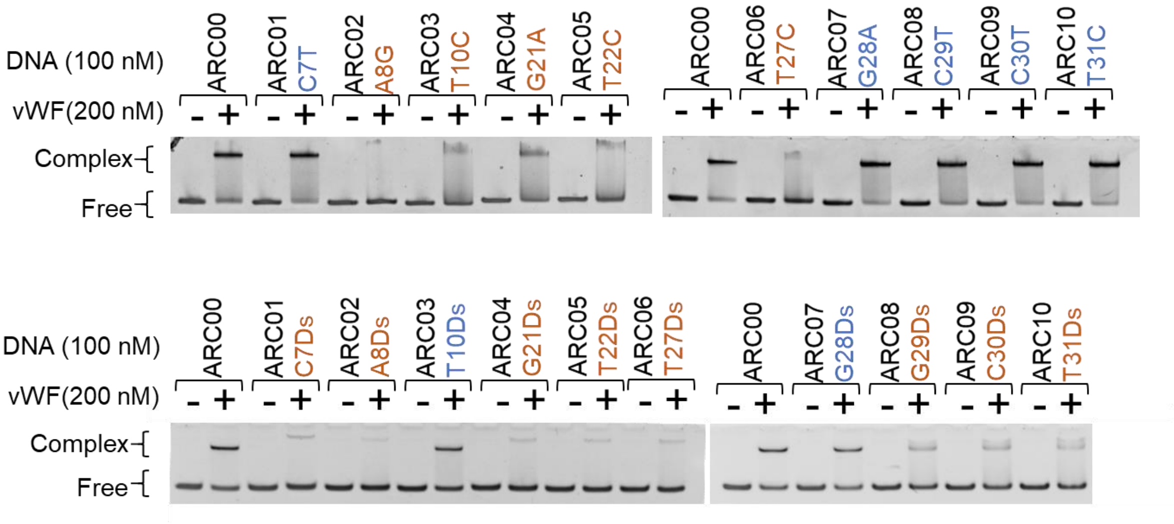

**Supplementary figure S9.** Hydrophobic binding sites detected near the aptamer–vWF binding interface. vWF is shown in white, and its binding partners are shown in yellow. (a) T10 corresponds to the binding site of Leu110 in botrocetin (PDB 1U0O). (b) The T31 site corresponds to the binding site of Tyr291 inbotrocetin (PDB 1U0O). (c) The T31 site corresponds to the binding site of Tyr316 in NMC-4 (PDB 1OAK).

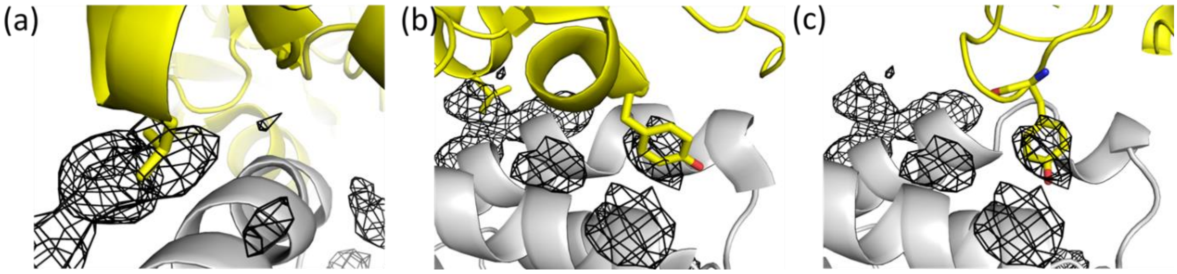


**Supplementary figure S10.** Binding modes of ARC1172 and its T27C variant with vWF. (a, b, c) Structural comparison of the ARC1172 and T27C variant complexes of vWF. (a) Crystal structure of vWF (white) complexed with ARC1172 (PDB 3HXQ), showing the hydrogen bonds formed by G26 and T27 with the backbone nitrogen of Arg1392 and the side chain of Gln1391, respectively. (b) Final MD trajectory structure of vWF complexed with ARC1172, showing the maintenance of the hydrogen bonds formed by G26 and T27 with the backbone nitrogen of Arg1392 and the side chain of Gln1391. (c) Final MD trajectory structure of vWF complexed with the T27C variant, showing the hydrogen bond between C27 and the side chain of Gln1391 and the loss of the hydrogen bond between G26 and Arg1392. (d) SPR sensorgrams for vWF binding with the stabilised T27C variant (top) and the original ARC1172 (bottom).

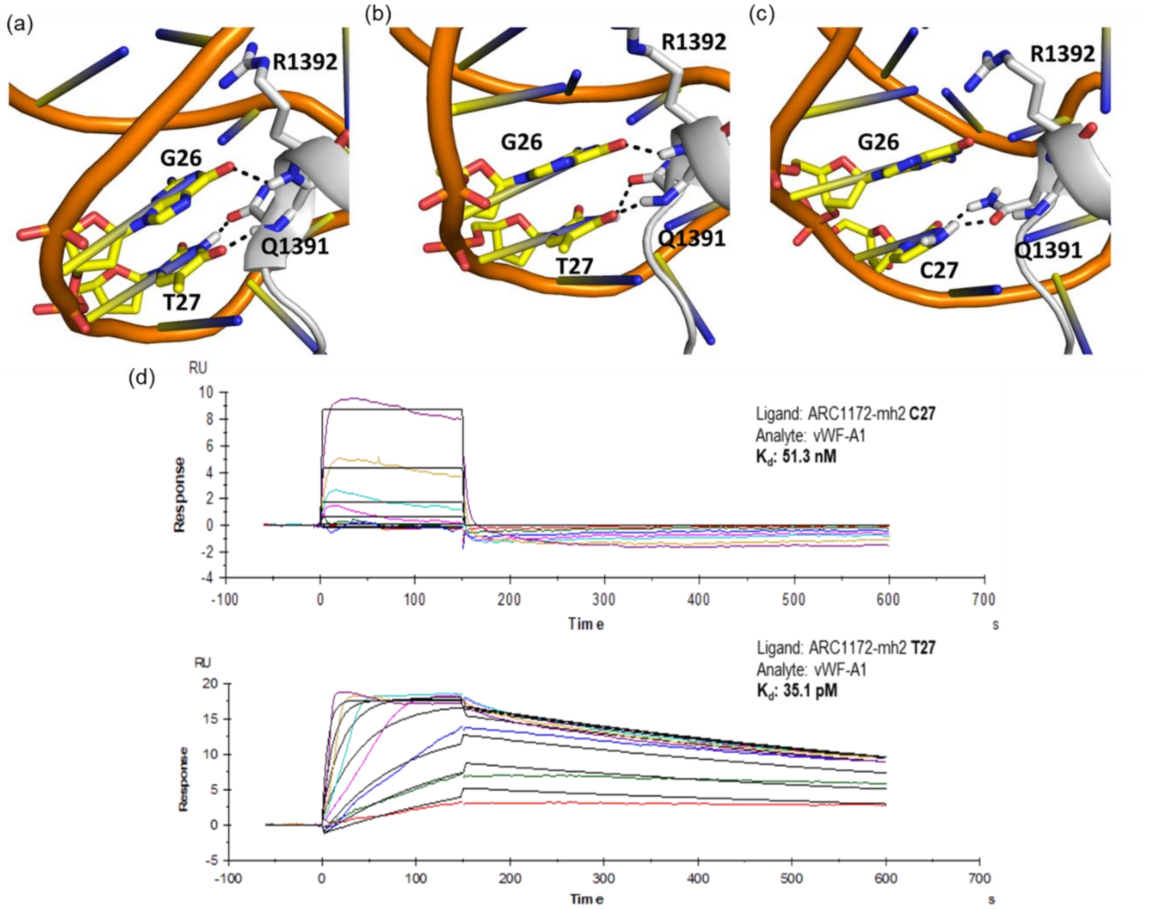

**Supplementary figure S11.** Cα root mean square deviations (RMSDs) for 10 independent ligand-mapping trajectories of vWF. Flexible N- and C-terminal residues were excluded from the calculations.

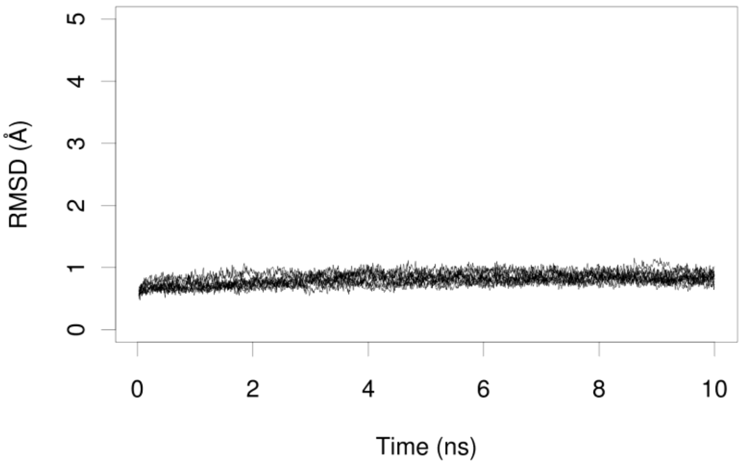

**Supplementary figure S12.** RMSDs for the Cα and DNA backbone atoms in the complexes of vWF with (A, B) ARC1172, (C, D) T10Ds, (E, F) G28Ds, and (G, H) T27C. Flexible N- and C-terminal vWF residues were excluded from the calculations.

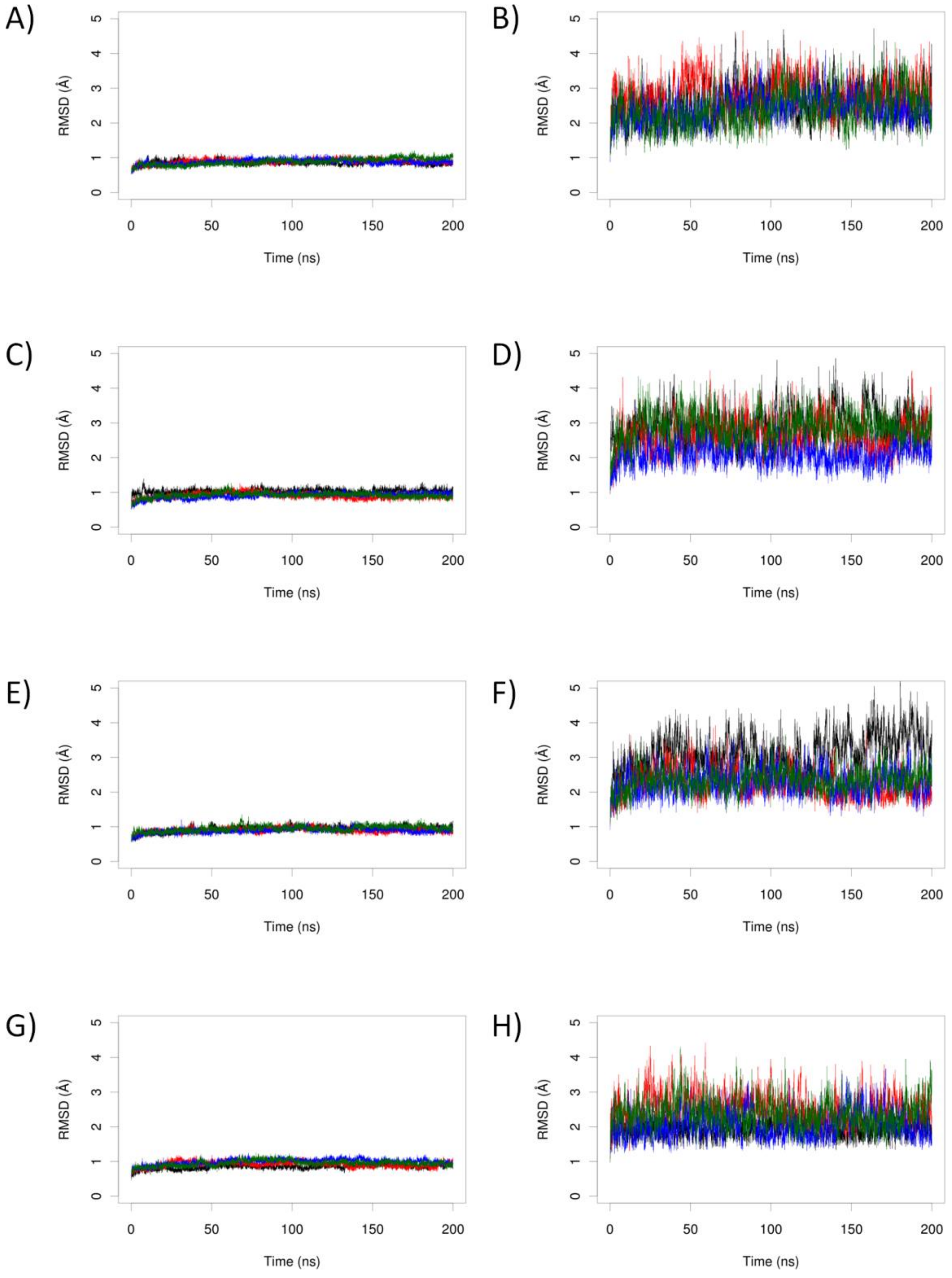

**Supplementary table S1**. Anti-IFNγ aptamer (48-mer) and its variants for the transition mutation experiments. The mini-hairpin DNA sequences are underlined. The mutated positions are indicated in bold and colored blue.

| Name | Original | Position (numbering) | Mutated | DNA sequences (5'- to -3'), I = Ds |
|---|---|---|---|---|
| IFN00 | - | - | - | CCCGCCCGGGTCCGCGAAGCGGTAGGTITGGGCTAG<br>GCIGCTGGCGGG |
| IFN01 | C | 7 | T | CCCGCC**T**GGGTCCGCGAAGCGGTAGGTITGGGCTAG<br>GCIGCTGGCGGG |
| IFN02 | G | 8 | A | CCCGCCC**A**GGTCCGCGAAGCGGTAGGTITGGGCTAG<br>GCIGCTGGCGGG |
| IFN03 | G | 9 | A | CCCGCCCG**A**GTCCGCGAAGCGGTAGGTITGGGCTAG<br>GCIGCTGGCGGG |
| IFN04 | G | 10 | A | CCCGCCCGG**A**TCCGCGAAGCGGTAGGTITGGGCTAG<br>GCIGCTGGCGGG |
| IFN05 | T | 11 | C | CCCGCCCGGG**C**CCGCGAAGCGGTAGGTITGGGCTAG<br>GCIGCTGGCGGG |
| IFN06 | T | 23 (24) | C | CCCGCCCGGGTCCGCGAAGCGG**C**AGGTITGGGCTAG<br>GCIGCTGGCGGG |
| IFN07 | A | 24 (25) | G | CCCGCCCGGGTCCGCGAAGCGGT**G**GGTITGGGCTAG<br>GCIGCTGGCGGG |
| IFN08 | G | 25 (26) | A | CCCGCCCGGGTCCGCGAAGCGGTA**A**GTITGGGCTAG<br>GCIGCTGGCGGG |
| IFN09 | G | 26 (27) | A | CCCGCCCGGGTCCGCGAAGCGGTAG**A**TITGGGCTAG<br>GCIGCTGGCGGG |
| IFN10 | T | 27 (28) | C | CCCGCCCGGGTCCGCGAAGCGGTAGG**C**ITGGGCTAG<br>GCIGCTGGCGGG |
| IFN11 | Ds | 28 (29) | A | CCCGCCCGGGTCCGCGAAGCGGTAGGT**A**TGGGCTAG<br>GCIGCTGGCGGG |
| IFN12 | T | 29 (30) | C | CCCGCCCGGGTCCGCGAAGCGGTAGGTI**C**GGGCTAG<br>GCIGCTGGCGGG |
| IFN13 | G | 30 (31) | A | CCCGCCCGGGTCCGCGAAGCGGTAGGTIT**A**GGCTAG<br>GCIGCTGGCGGG |
| IFN14 | G | 31 (32) | A | CCCGCCCGGGTCCGCGAAGCGGTAGGTITG**A**GCTAG<br>GCIGCTGGCGGG |
| IFN15 | G | 32 (33) | A | CCCGCCCGGGTCCGCGAAGCGGTAGGTITGG**A**CTAG<br>GCIGCTGGCGGG |
| IFN16 | C | 33 (34) | T | CCCGCCCGGGTCCGCGAAGCGGTAGGTITGGG**T**TAG<br>GCIGCTGGCGGG |
| IFN17 | T | 34 (35) | C | CCCGCCCGGGTCCGCGAAGCGGTAGGTITGGGC**C**AG<br>GCIGCTGGCGGG |
| IFN18 | A | 35 (36) | G | CCCGCCCGGGTCCGCGAAGCGGTAGGTITGGGCT**G**G<br>GCIGCTGGCGGG |
| IFN19 | G | 36 (37) | A | CCCGCCCGGGTCCGCGAAGCGGTAGGTITGGGCTA**A**<br>GCIGCTGGCGGG |
| IFN20 | G | 37 (38) | A | CCCGCCCGGGTCCGCGAAGCGGTAGGTITGGGCTAG<br>**A**CIGCTGGCGGG |
| IFN21 | C | 38 (39) | T | CCCGCCCGGGTCCGCGAAGCGGTAGGTITGGGCTAG<br>G**T**IGCTGGCGGG |
| IFN22 | Ds | 39 (40) | A | CCCGCCCGGGTCCGCGAAGCGGTAGGTITGGGCTAG<br>GC**A**GCTGGCGGG |
| IFN23 | G | 40 (41) | A | CCCGCCCGGGTCCGCGAAGCGGTAGGTITGGGCTAG<br>GCI**A**CTGGCGGG |
| IFN24 | C | 41 (42) | T | CCCGCCCGGGTCCGCGAAGCGGTAGGTITGGGCTAG<br>GCIG**T**TGGCGGG |
| IFN25 | T | 42 (43) | C | CCCGCCCGGGTCCGCGAAGCGGTAGGTITGGGCTAG<br>GCIGC**C**GGCGGG |

**Supplementary table S2**. Anti-vWF aptamer (42-mer) and its variants for the transition mutation experiments. The mini-hairpin DNA sequences are underlined. The mutated positions are indicated in bold.

| Name | Original | Position (numbering) | Mutated | DNA sequences (5'- to -3'),<br>I = Ds |
|---|---|---|---|---|
| vWF00 | - | - | - | CGCGGCCGAIGAGCACCGAAGGTCTCAACGITGG<br>AGGCCGCG |
| vWF01 | G | 8 | A | CGCGGCC**A**AIGAGCACCGAAGGTCTCAACGITGG<br>AGGCCGCG |
| vWF02 | A | 9 | G | CGCGGCCG**G**IGAGCACCGAAGGTCTCAACGITGG<br>AGGCCGCG |
| vWF03 | Ds | 10 | A | CGCGGCCGA**A**GAGCACCGAAGGTCTCAACGITGG<br>AGGCCGCG |
| vWF04 | C | 14 | T | CGCGGCCGAIGAG**T**ACCGAAGGTCTCAACGITGG<br>AGGCCGCG |
| vWF05 | A | 27 (29) | G | CGCGGCCGAIGAGCACCGAAGGTCTC**G**ACGITGG<br>AGGCCGCG |
| vWF06 | A | 28 (30) | G | CGCGGCCGAIGAGCACCGAAGGTCTCA**G**CGITGG<br>AGGCCGCG |
| vWF07 | C | 29 (31) | T | CGCGGCCGAIGAGCACCGAAGGTCTCAA**T**GITGG<br>AGGCCGCG |
| vWF08 | G | 30 (32) | A | CGCGGCCGAIGAGCACCGAAGGTCTCAAC**A**ITGG<br>AGGCCGCG |
| vWF09 | Ds | 31 (33) | A | CGCGGCCGAIGAGCACCGAAGGTCTCAACG**A**TGG<br>AGGCCGCG |
| vWF10 | T | 32 (34) | C | CGCGGCCGAIGAGCACCGAAGGTCTCAACGI**C**GG<br>AGGCCGCG |
| vWF11 | G | 33 (35) | A | CGCGGCCGAIGAGCACCGAAGGTCTCAACGIT**A**G<br>AGGCCGCG |
| vWF12 | G | 34 (36) | A | CGCGGCCGAIGAGCACCGAAGGTCTCAACGITG**A**<br>AGGCCGCG |
| vWF13 | A | 35 (37) | G | CGCGGCCGAIGAGCACCGAAGGTCTCAACGITGG<br>**G**GGCCGCG |

**Supplementary table S3**. Anti-DENV1-NS1 aptamer AptD1 (48-mer) and its variants for the transition mutation experiments. The mini-hairpin DNA sequences are underlined. The mutated positions are indicated in bold.

| Name | Original | Position (numbering) | Mutated | DNA sequences (5'- to -3'),<br>I = Ds |
|---|---|---|---|---|
| AptD1-00 | - | - | - | CCCCAGACGGACTGGTGTICTCGGIATGGCC<br>GTCTGGGGCGCGAAGCG |
| AptD1-01 | A | 11 (7) | G | CCCCAGACGG**G**CTGGTGTICTCGGIATGGCC<br>GTCTGGGGCGCGAAGCG |
| AptD1-02 | C | 12 (8) | T | CCCCAGACGGA**T**TGGTGTICTCGGIATGGCC<br>GTCTGGGGCGCGAAGCG |
| AptD1-03 | T | 13 (9) | C | CCCCAGACGGAC**C**GGTGTICTCGGIATGGCC<br>GTCTGGGGCGCGAAGCG |
| AptD1-04 | G | 14 (10) | A | CCCCAGACGGACT**A**GTGTICTCGGIATGGCC<br>GTCTGGGGCGCGAAGCG |
| AptD1-05 | G | 15 (11) | A | CCCCAGACGGACTG**A**TGTICTCGGIATGGCC<br>GTCTGGGGCGCGAAGCG |
| AptD1-06 | T | 16 (12) | C | CCCCAGACGGACTGG**C**GTICTCGGIATGGCC<br>GTCTGGGGCGCGAAGCG |
| AptD1-07 | G | 17 (13) | A | CCCCAGACGGACTGGT**A**TICTCGGIATGGCC<br>GTCTGGGGCGCGAAGCG |
| AptD1-08 | T | 18 (14) | C | CCCCAGACGGACTGGTG**C**ICTCGGIATGGCC<br>GTCTGGGGCGCGAAGCG |
| AptD1-09 | Ds | 19 (15) | A | CCCCAGACGGACTGGTGT**A**CTCGGIATGGCC<br>GTCTGGGGCGCGAAGCG |
| AptD1-10 | C | 20 (16) | T | CCCCAGACGGACTGGTGTI**T**TCGGIATGGCC<br>GTCTGGGGCGCGAAGCG |
| AptD1-11 | T | 21 (17) | C | CCCCAGACGGACTGGTGTIC**C**CGGIATGGCC<br>GTCTGGGGCGCGAAGCG |
| AptD1-12 | C | 22 (18) | T | CCCCAGACGGACTGGTGTICT**T**GGIATGGCC<br>GTCTGGGGCGCGAAGCG |
| AptD1-13 | G | 23 (19) | A | CCCCAGACGGACTGGTGTICTC**A**GIATGGCC<br>GTCTGGGGCGCGAAGCG |
| AptD1-14 | G | 24 (20) | A | CCCCAGACGGACTGGTGTICTCG**A**IATGGCC<br>GTCTGGGGCGCGAAGCG |
| AptD1-15 | Ds | 25 (21) | A | CCCCAGACGGACTGGTGTICTCGG**A**ATGGCC<br>GTCTGGGGCGCGAAGCG |
| AptD1-16 | A | 26 (22) | G | CCCCAGACGGACTGGTGTICTCGGI**G**TGGCC<br>GTCTGGGGCGCGAAGCG |
| AptD1-17 | T | 27 (23) | C | CCCCAGACGGACTGGTGTICTCGGIA**C**GGCC<br>GTCTGGGGCGCGAAGCG |
| AptD1-18 | G | 28 (24) | A | CCCCAGACGGACTGGTGTICTCGGIAT**A**GCC<br>GTCTGGGGCGCGAAGCG |
| AptD1-19 | G | 29 (25) | A | CCCCAGACGGACTGGTGTICTCGGIATG**A**CC<br>GTCTGGGGCGCGAAGCG |

**Supplementary table S4**. Anti-DENV3-NS1 aptamer (50-mer) and its variants for the transition mutation experiments. The mutated positions are indicated in bold.

| Name | Original | Position (numbering) | Mutated | DNA sequences (5'- to -3'), I = Ds |
|---|---|---|---|---|
| AptD3-00 | - | - | - | CCGCTTGTCATCTAICCTGGCCITGTGGTAC TGTAACGGCTGACAAGCGG |
| AptD3-01 | T | 11 (7) | C | CCGCTTGTCA**C**CTAICCTGGCCITGTGGTAC TGTAACGGCTGACAAGCGG |
| AptD3-02 | C | 12 (8) | T | CCGCTTGTCAT**T**TAICCTGGCCITGTGGTAC TGTAACGGCTGACAAGCGG |
| AptD3-03 | T | 13 (9) | C | CCGCTTGTCATC**C**AICCTGGCCITGTGGTAC TGTAACGGCTGACAAGCGG |
| AptD3-04 | A | 14 (10) | G | CCGCTTGTCATCT**G**ICCTGGCCITGTGGTAC TGTAACGGCTGACAAGCGG |
| AptD3-05 | Ds | 15 (11) | A | CCGCTTGTCATCTA**A**CCTGGCCITGTGGTAC TGTAACGGCTGACAAGCGG |
| AptD3-06 | C | 16 (12) | T | CCGCTTGTCATCTAI**T**CTGGCCITGTGGTAC TGTAACGGCTGACAAGCGG |
| AptD3-07 | C | 17 (13) | T | CCGCTTGTCATCTAIC**T**TGGCCITGTGGTAC TGTAACGGCTGACAAGCGG |
| AptD3-08 | T | 18 (14) | C | CCGCTTGTCATCTAICC**C**GGCCITGTGGTAC TGTAACGGCTGACAAGCGG |
| AptD3-09 | G | 19 (15) | A | CCGCTTGTCATCTAICCT**A**GCCITGTGGTAC TGTAACGGCTGACAAGCGG |
| AptD3-10 | G | 20 (16) | A | CCGCTTGTCATCTAICCTG**A**CCITGTGGTAC TGTAACGGCTGACAAGCGG |
| AptD3-11 | C | 21 (17) | T | CCGCTTGTCATCTAICCTGG**T**CITGTGGTAC TGTAACGGCTGACAAGCGG |
| AptD3-12 | C | 22 (18) | T | CCGCTTGTCATCTAICCTGGC**T**ITGTGGTAC TGTAACGGCTGACAAGCGG |
| AptD3-13 | Ds | 23 (19) | A | CCGCTTGTCATCTAICCTGGCC**A**TGTGGTAC TGTAACGGCTGACAAGCGG |
| AptD3-14 | T | 24 (20) | C | CCGCTTGTCATCTAICCTGGCCI**C**GTGGTAC TGTAACGGCTGACAAGCGG |
| AptD3-15 | G | 25 (21) | A | CCGCTTGTCATCTAICCTGGCCIT**A**TGGTAC TGTAACGGCTGACAAGCGG |
| AptD3-16 | T | 26 (22) | C | CCGCTTGTCATCTAICCTGGCCITG**C**GGTAC TGTAACGGCTGACAAGCGG |
| AptD3-17 | G | 27 (23) | A | CCGCTTGTCATCTAICCTGGCCITGT**A**GTAC TGTAACGGCTGACAAGCGG |
| AptD3-18 | G | 28 (24) | A | CCGCTTGTCATCTAICCTGGCCITGTG**A**TAC TGTAACGGCTGACAAGCGG |
| AptD3-19 | T | 29 (25) | C | CCGCTTGTCATCTAICCTGGCCITGTGG**C**AC TGTAACGGCTGACAAGCGG |
| AptD3-20 | A | 30 (26) | G | CCGCTTGTCATCTAICCTGGCCITGTGGT**G**C TGTAACGGCTGACAAGCGG |
| AptD3-21 | C | 31 (27) | T | CCGCTTGTCATCTAICCTGGCCITGTGGTA**T** TGTAACGGCTGACAAGCGG |
| AptD3-22 | T | 32 (28) | C | CCGCTTGTCATCTAICCTGGCCITGTGGTAC **C**GTAACGGCTGACAAGCGG |
| AptD3-23 | G | 33 (29) | A | CCGCTTGTCATCTAICCTGGCCITGTGGTAC T**A**TAACGGCTGACAAGCGG |
| AptD3-24 | T | 34 (30) | C | CCGCTTGTCATCTAICCTGGCCITGTGGTAC TG**C**AACGGCTGACAAGCGG |
| AptD3-25 | A | 35 (31) | G | CCGCTTGTCATCTAICCTGGCCITGTGGTAC TGT**G**ACGGCTGACAAGCGG |

| | | | | |
|---|---|---|---|---|
| AptD3-26 | A | 36 (32) | G | CCGCTTGTCATCTAICCTGGCCITGTGGTAC TGTA**G**CGGCTGACAAGCGG |
| AptD3-27 | C | 37 (33) | T | CCGCTTGTCATCTAICCTGGCCITGTGGTAC TGTAA**T**GGCTGACAAGCGG |
| AptD3-28 | G | 38 (34) | A | CCGCTTGTCATCTAICCTGGCCITGTGGTAC TGTAAC**A**GCTGACAAGCGG |
| AptD3-29 | G | 39 (35) | A | CCGCTTGTCATCTAICCTGGCCITGTGGTAC TGTAACG**A**CTGACAAGCGG |
| AptD3-30 | C | 40 (36) | T | CCGCTTGTCATCTAICCTGGCCITGTGGTAC TGTAACGG**T**TGACAAGCGG |

**Supplementary table S5**. Anti-vWF aptamer ARC1172 (41-mer) and its variants for the transition mutation and Ds-replacement experiments. The mutated positions are indicated in bold.

| Name | Original | Position (numbering) | Mutated | DNA sequences (5'- to -3'),<br>I = Ds |
|---|---|---|---|---|
| ARC00 | - | - | - | GGCGTGCAGTGCCTTCGGCCGTGCGGTGCCT<br>CCGTCACGCC |
| ARC01 | C | 7 | T | GGCGTG**T**AGTGCCTTCGGCCGTGCGGTGCCT<br>CCGTCACGCC |
| ARC02 | A | 8 | G | GGCGTGC**G**GTGCCTTCGGCCGTGCGGTGCCT<br>CCGTCACGCC |
| ARC03 | T | 10 | C | GGCGTGCAG**C**GCCTTCGGCCGTGCGGTGCCT<br>CCGTCACGCC |
| ARC04 | G | 21 | A | GGCGTGCAGTGCCTTCGGCC**A**TGCGGTGCCT<br>CCGTCACGCC |
| ARC05 | T | 22 | C | GGCGTGCAGTGCCTTCGGCCG**C**GCGGTGCCT<br>CCGTCACGCC |
| ARC06 | T | 27 | C | GGCGTGCAGTGCCTTCGGCCGTGCGG**C**GCCT<br>CCGTCACGCC |
| ARC07 | G | 28 | A | GGCGTGCAGTGCCTTCGGCCGTGCGGT**A**CCT<br>CCGTCACGCC |
| ARC08 | C | 29 | T | GGCGTGCAGTGCCTTCGGCCGTGCGGTG**T**CT<br>CCGTCACGCC |
| ARC09 | C | 30 | T | GGCGTGCAGTGCCTTCGGCCGTGCGGTGC**T**T<br>CCGTCACGCC |
| ARC10 | T | 31 | C | GGCGTGCAGTGCCTTCGGCCGTGCGGTGCC**C**<br>CCGTCACGCC |
| ARC01Ds | C | 7 | Ds | GGCGTG**I**AGTGCCTTCGGCCGTGCGGTGCCT<br>CCGTCACGCC |
| ARC02Ds | A | 8 | Ds | GGCGTGC**I**GTGCCTTCGGCCGTGCGGTGCCT<br>CCGTCACGCC |
| ARC03Ds | T | 10 | Ds | GGCGTGCAG**I**GCCTTCGGCCGTGCGGTGCCT<br>CCGTCACGCC |
| ARC04Ds | G | 21 | Ds | GGCGTGCAGTGCCTTCGGCC**I**TGCGGTGCCT<br>CCGTCACGCC |
| ARC05Ds | T | 22 | Ds | GGCGTGCAGTGCCTTCGGCCG**I**GCGGTGCCT<br>CCGTCACGCC |
| ARC06Ds | T | 27 | Ds | GGCGTGCAGTGCCTTCGGCCGTGCGG**I**GCCT<br>CCGTCACGCC |
| ARC07Ds | G | 28 | Ds | GGCGTGCAGTGCCTTCGGCCGTGCGGT**I**CCT<br>CCGTCACGCC |
| ARC08Ds | C | 29 | Ds | GGCGTGCAGTGCCTTCGGCCGTGCGGTG**I**CT<br>CCGTCACGCC |
| ARC09Ds | C | 30 | Ds | GGCGTGCAGTGCCTTCGGCCGTGCGGTGC**I**T<br>CCGTCACGCC |
| ARC10Ds | T | 31 | Ds | GGCGTGCAGTGCCTTCGGCCGTGCGGTGCC**I**<br>CCGTCACGCC |

**Supplementary table S6**. Experimental conditions for EMSA.

| Aptamer variants | Target | Binding Buffer | Gel (with 5% glycerol) | Running buffer |
|---|---|---|---|---|
| IFN series: 25 nM | IFNγ : 50 nM | 1× PBS, 0.05% Nonidet P-40 | 8% polyacrylamide (29:1), 0.5× TBE, 3 M Urea | 0.5× TBE |
| vWF series: 25 nM | vWF: 50 nM | | | |
| AptD1 series: 50 nM | DENV1 NS1: 25 nM (hexamer) | 20 mM Tris-HCl, pH 7.5, 150 mM NaCl, 1 mM $MgCl_2$, 2.7 mM KCl | 4% polyacrylamide (29:1), 0.5× TB, 1 mM $MgCl_2$, 2.7 mM KCl | 0.5× TB, 1 mM $MgCl_2$, 2.7 mM KCl |
| AptD3 series: 50 nM | DENV3 NS1: 25 nM (hexamer) | | | |
| ARC series: 100 nM | vWF: 200 nM | 1× PBS, 0.05% Nonidet P-40 | 10% polyacrylamide (29:1), 0.5× TBE | 0.5× TBE |

**Supplementary table S7**. Computed average binding free energy of ARC1172 aptamer and its variants.

| Aptamer | Sequence | ΔG (kcal/mol) |
|---|---|---|
| ARC1172 | GGCGTGCAG**T**GCCTTCGGCCGTGCGGT**G**CCTCCGTCACGCC | -188.3 ± 3.1 |
| T10Ds | GGCGTGCAG**Ds**GCCTTCGGCCGTGCGGT**G**CCTCCGTCACGCC | -192.7 ± 4.3 |
| G28Ds | GGCGTGCAG**T**GCCTTCGGCCGTGCGGT**Ds**CCTCCGTCACGCC | -193.9 ± 3.8 |
| T27C | GGCGTGCAG**T**GCCTTCGGCCGTGCGG**CG**CCTCCGTCACGCC | -174.7 ± 9.7 |

### Methods

#### Surface Plasmon Resonance (SPR) analysis

The SPR analysis was performed to compare the vWF binding affinity profiles of ARC1172 aptamer variants with the internal and 3'-terminal mini-hairpin structures (ARC1172-mh2; 5'-GGCGTGCAGTGCCGAAGGCCGTGCGG**N**GCCTCCGTCACGCCCGCG(Biotin-dT)AGCG-3', the mini-hairpin sequences are underlined and **N** (T or C) is in bold font), using a Biacore T200 (GE Healthcare) platform. Each biotinylated aptamer variant (ligand) was immobilised on the flow cells of streptavidin-coated sensor chips, by injecting 0.5 nM of ligand solution in running buffer (1× PBS with 0.05% Nonidet P-40) at a flow rate of 5 µL/min for 240 or 960 sec (respectively for **N** = C or T) at 25°C. Analyte (vWF) solutions from 0.078125 nM to 5 nM were prepared in running buffer and injected at a flow rate of 100 µL/min, with an association duration of 150 sec and a dissociation duration of 450 sec. The sensor chip was regenerated with a 5-sec injection of 50 mM NaOH, followed by a 10-min equilibration with running buffer. Using the BIAevaluation T200 software version 1.0 (GE Healthcare), the $K_D$ values were determined by fitting the blank-subtracted kinetics data with a 1:1 binding model.

#### Molecular dynamics simulations

##### Preparation of structures

The crystal structures of the A1 domain of von Willebrand factor (vWF) bound to the DNA aptamer ARC1172 (PDB code 3HXQ[1]) and the apo A1 domain of vWF (PDB code 1AUQ[2]) were used as the starting structures for conventional molecular dynamics (MD) and ligand-mapping molecular dynamics (LMMD) simulations, respectively. Complexes of vWF with ARC1172, T10Ds, G28Ds, and T27C variants (Supplementary table S7) were prepared for MD simulations. The N- and C-termini of vWF were capped by an acetyl group and an N-methyl group, respectively. PyMOL[3] was used to convert Thy10 and Gua28 to Ds and Thy27 to Cyt in ARC1172. The protonation states of residues were determined with the PDB2PQR.program.[4] Each system was then solvated with TIP3P water molecules[5] in a periodic truncated octahedron box with its walls at least 10 Å away from the protein and/or aptamer, followed by charge neutralisation with sodium ions.

##### Molecular dynamics (MD)

Four independent MD simulations were performed for each of the vWF–aptamer complexes. Energy minimisation and MD simulation were performed with the PMEMD module of AMBER 18[6] using the ff14SB[7] force field for the protein and the OL15[8] force field for DNA. The Ds nucleotide was described by both the OL15 and generalised AMBER force fields (GAFF).[9] Atomic charges for Ds were derived using the R.E.D. Server[10] by fitting restrained electrostatic potential (RESP) charges[11] to a molecular electrostatic potential computed by the Gaussian 16 program[12] at the HF/6-31G* level of theory. A time step of 2 fs was used and the SHAKE algorithm[13] was implemented to constrain all bonds involving hydrogen atoms. The particle mesh Ewald method[14] was used to treat long-range electrostatic interactions under periodic boundary conditions. A cutoff distance of 9 Å was implemented for nonbonded interactions. The non-hydrogen atoms of the protein and aptamer were maintained with a harmonic positional restraint of 2.0 kcal $mol^{-1}$ $Å^{-2}$ during the minimisation and equilibration steps. Energy minimisation was performed using the steepest descent algorithm for 500 steps, followed by another 500 steps with the conjugate gradient algorithm. Gradual heating of the systems to 300 K was performed at constant volume over 50 ps before equilibration at a constant pressure of 1 atm for another 50 ps. The restraints were removed for the subsequent equilibration (2 ns) and production (200 ns) runs, which were performed at 300 K using a Langevin thermostat[15] with a collision frequency of 2 $ps^{-1}$, and at 1 atm using a Berendsen barostat[16] with a pressure relaxation time of 2 ps.

##### Ligand-mapping molecular dynamics (LMMD)

LMMD simulations were performed on the A1 domain of vWF. Ten different distributions of benzenes around the protein were created, using Packmol.[17] The LEaP module in the AMBER 18 package was then used to solvate each system with TIP3P water molecules in a periodic truncated octahedron box, such that its walls were at least 10 Å away from the protein, and for neutralisation of charges with chloride ions, resulting in a final benzene concentration of ~0.2 M. Minimisation, equilibration and production (10 ns) MD simulations were performed as described above, for a cumulative sampling time of 100 ns. The GAFF[9] force field was used to describe the benzenes during the simulations. Atomic charges for benzene (Table S3) were obtained from a previous report.[18]

**Binding free energy calculations**

Binding free energies for the vWF–aptamer complexes were calculated using the molecular mechanics/generalised Born surface area (MM/GBSA) method[19] implemented in AMBER 14.[20] Two hundred equally-spaced snapshot structures were extracted from the last 60 (the T27C variant), 80 (the T10Ds variant) or 100 ns (the G28Ds variant) of each of the trajectories, and their molecular mechanical energies were calculated with the sander module. The polar contribution to the solvation free energy was calculated by the pbsa[21] program using the modified generalised Born (GB) model described by Onufriev *et al.*[22], with the solute dielectric constant set to 4 and the exterior dielectric constant set to 80. The nonpolar contribution was estimated from the solvent accessible surface area using the molsurf[23] program, with $\gamma$ = 0.005 kcal Å$^{-2}$ and $\beta = 0$. The entropy contribution was not considered, as it has been shown to be unnecessary for ranking the binding affinities of structurally similar ligands.[24]


(1) Huang, R.-H.; Fremont, D. H.; Diener, J. L.; Schaub, R. G.; Sadler, J. E., A structural explanation for the antithrombotic activity of ARC1172, a DNA aptamer that binds von Willebrand factor domain A1. *Structure* **2009,** *17*, 1476-1484.
(2) Emsley, J.; Cruz, M.; Handin, R.; Liddington, R., Crystal structure of the von Willebrand Factor A1 domain and implications for the binding of platelet glycoprotein Ib. *J. Biol. Chem.* **1998,** *273*, 10396-10401.
(3) DeLano, W. L. *The PyMOL Molecular Graphics System*; DeLano Scientific: San Carlos, CA, USA, 2002.
(4) Dolinsky, T. J.; Czodrowski, P.; Li, H.; Nielsen, J. E.; Jensen, J. H.; Klebe, G.; Baker, N. A., PDB2PQR: expanding and upgrading automated preparation of biomolecular structures for molecular simulations. *Nucleic Acids Res.* **2007,** *35*, W522-W525.
(5) Jorgensen, W. L.; Chandrasekhar, J.; Madura, J. D.; Impey, R. W.; Klein, M. L., Comparison of simple potential functions for simulating liquid water. *J. Chem. Phys.* **1983,** *79*, 926-935.
(6) Case, D. A.; Ben-Shalom, I. Y.; Brozell, S. R.; Cerutti, D. S.; Cheatham, T. E., III; Cruzeiro, V. W. D.; Darden, T. A.; Duke, R. E.; Ghoreishi, D.; Gilson, M. K.; Gohlke, H.; Goetz, A. W.; Greene, D.; Harris, R.; Homeyer, N.; Izadi, S.; Kovalenko, A.; Kurtzman, T.; Lee, T. S.; LeGrand, S.; Li, P.; Lin, C.; Liu, J.; Luchko, T.; Luo, R.; Mermelstein, D. J.; Merz, K. M.; Miao, Y.; Monard, G.; Nguygen, C.; Nguygen, H.; Omelyan, I.; Onufriev, A.; Pan, F.; Qi, R.; Roe, D. R.; Roitberg, A.; Sagui, C.; Schott-Verdugo, S.; Shen, J.; Simmerling, C. L.; Smith, J.; Salomon-Ferrer, R.; Swails, J.; Walker, R. C.; Wang, J.; Wei, H.; Wolf, R. M.; Wu, X.; Xiao, L.; York, D. M.; Kollman, P. A. *AMBER 18*, University of California, San Francisco: 2018.
(7) Maier, J. A.; Martinez, C.; Kasavajhala, K.; Wickstrom, L.; Hauser, K. E.; Simmerling, C., ff14SB: improving the accuracy of protein side chain and backbone parameters from ff99SB. *J. Chem. Theory Comput.* **2015,** *11*, 3696-3713.
(8) Zgarbová, M.; Šponer, J.; Otyepka, M.; Cheatham, T. E.; Galindo-Murillo, R.; Jurečka, P., Refinement of the sugar–phosphate backbone torsion beta for AMBER force fields improves the description of Z- and B-DNA. *J. Chem. Theory Comput.* **2015,** *11*, 5723-5736.
(9) Wang, J. M.; Wolf, R. M.; Caldwell, J. W.; Kollman, P. A.; Case, D. A., Development and testing of a general amber force field. *J. Comput. Chem.* **2004,** *25*, 1157-1174.
(10) Vanquelef, E.; Simon, S.; Marquant, G.; Garcia, E.; Klimerak, G.; Delepine, J. C.; Cieplak, P.; Dupradeau, F.-Y., R.E.D. Server: a web service for deriving RESP and ESP charges and building force field libraries for new molecules and molecular fragments. *Nucleic Acids Res.* **2011,** *39*, W511-W517.
(11) Cornell, W. D.; Cieplak, P.; Bayly, C. I.; Kollman, P. A., Application of RESP charges to calculate conformational energies, hydrogen bond energies, and free energies of solvation. *J. Am. Chem. Soc.* **1993,** *115*, 9620-9631.
(12) Frisch, M. J.; Trucks, G. W.; Schlegel, H. B.; Scuseria, G. E.; Robb, M. A.; Cheeseman, J. R.; Scalmani, G.; Barone, V.; Petersson, G. A.; Nakatsuji, H.; Li, X.; Caricato, M.; Marenich, A. V.; Bloino, J.; Janesko, B. G.; Gomperts, R.; Mennucci, B.; Hratchian, H. P.; Ortiz, J. V.; Izmaylov, A. F.; Sonnenberg, J. L.; Williams-Young, D.; Ding, F.; Lipparini, F.; Egidi, F.; Goings, J.; Peng, B.; Petrone, A.; Henderson, T.; Ranasinghe, D.; Zakrzewski, V. G.; Gao, J.; Rega, N.; Zheng, G.; Liang, W.; Hada, M.; Ehara, M.; Toyota, K.; Fukuda, R.; Hasegawa, J.; Ishida, M.; Nakajima, T.; Honda, Y.; Kitao, O.; Nakai, H.; Vreven, T.; Throssell, K.; Montgomery Jr., J. A.; Peralta, J. E.; Ogliaro, F.; Bearpark, M. J.; Heyd, J. J.; Brothers, E. N.;

Kudin, K. N.; Staroverov, V. N.; Keith, T. A.; Kobayashi, R.; Normand, J.; Raghavachari, K.; Rendell, A. P.; Burant, J. C.; Iyengar, S. S.; Tomasi, J.; Cossi, M.; Millam, J. M.; Klene, M.; Adamo, C.; Cammi, R.; Ochterski, J. W.; Martin, R. L.; Morokuma, K.; Farkas, O.; Foresman, J. B.; Fox, D. J. *Gaussian 16 Rev. C.01*, Wallingford, CT, 2016.

(20) Case, D. A.; Babin, V.; Berryman, J. T.; Betz, R. M.; Cai, Q.; Cerutti, D. S.; Cheatham, T. E., III; Darden, T. A.; Duke, R. E.; Gohlke, H.; Goetz, A. W.; Gusarov, S.; Homeyer, N.; Janowski, P.; Kaus, J.; Kolossváry, I.; Kovalenko, A.; Lee, T. S.; LeGrand, S.; Luchko, T.; Luo, R.; Madej, B.; Merz, K. M.; Paesani, F.; Roe, D. R.; Roitberg, A.; Sagui, C.; Salomon-Ferrer, R.; Seabra, G.; Simmerling, C. L.; Smith, W.; Swails, J.; Walker, R. C.; Wang, J.; Wolf, R. M.; Wu, X.; Kollman, P. A. *AMBER 14*, University of California, San Francisco: 2014.

(21) Luo, R.; David, L.; Gilson, M. K., Accelerated Poisson-Boltzmann calculations for static and dynamic systems. *J. Comput. Chem.* **2002,** *23*, 1244-1253.

(22) Onufriev, A.; Bashford, D.; Case, D. A., Exploring protein native states and large-scale conformational changes with a modified generalized Born model. *Proteins: Struct. Funct. Bioinform.* **2004,** *55*, 383-394.

(23) Connolly, M. L., Analytical molecular surface calculation. *J. Appl. Crystallogr.* **1983,** *16*, 548-558.

(24) Hou, T.; Wang, J.; Li, Y.; Wang, W., Assessing the performance of the MM/PBSA and MM/GBSA methods. 1. The accuracy of binding free energy calculations based on molecular dynamics simulations. *J. Chem. Inf. Model.* **2011,** *51*, 69-82.
